# The Molecules Gateway: a homogeneous, searchable database of 150k annotated molecules from Actinomycetes

**DOI:** 10.1101/2024.06.28.601135

**Authors:** Matteo Simone, Marianna Iorio, Paolo Monciardini, Massimo Santini, Niccolò Cantù, Arianna Tocchetti, Stefania Serina, Cristina Brunati, Thomas Vernay, Andrea Gentile, Mattia Aracne, Marco Cozzi, Justin J.J. van der Hooft, Margherita Sosio, Stefano Donadio, Sonia I. Maffioli

**Affiliations:** NAICONS SRL, Milan, Italy; University of Milan, Milan, Italy; Code Atlas SRL, Legnano, Italy; University of Milano-Bicocca, Milan; Wageningen University, Wageningen, The Netherlands

**Author notes:** These authors contributed equally (Matteo Simone, Marianna Iorio, Paolo Monciardini).

## Abstract

Natural products are a sustainable resource for drug discovery, but their identification in complex mixtures remains a daunting task. We present an automated pipeline that compares, harmonizes and ranks the annotations of LC-HRMS data by different tools. When applied to 7,400 extracts derived from 6,566 strains belonging to 86 actinomycete genera, it yielded 150,000 molecules after processing over 50 million MS features. The web-based Molecules Gateway provides a highly interactive access to experimental and calculated data for these molecules, along with the metadata related to extracts and producer strains. We show how the Molecules Gateway can be used to rapidly identify known hard to find microbial products, unreported analogs of known families and not yet described metabolites. The Molecules Gateway, which complements available repositories of annotated MS data, is experimentally and computationally homogeneous, and thus amenable to global analyses, which show a large and untapped chemical diversity afforded by actinomycetes.

---

Natural products (NPs), molecules produced by living organisms, continue to provide a longstanding source of diverse drugs and drug leads^1^. The entire chemical complexity of an organism, which can consist of a myriad of molecules, can be captured by processing the entire organism, or portion(s) thereof, with appropriate methods to prepare extracts, each containing many molecules of unknown nature and concentration. While properly prepared extracts are suitable for most drug discovery programs, the identification of the molecule(s) responsible for the observed biological signal requires the time-consuming processes of deconvolution, i.e., reducing the number of molecules present in the sample to be tested, and dereplication, i.e., matching the observed properties of the active molecule(s) to those reported in available databases^2^. Hence, knowing *a priori* the composition of extracts would greatly accelerate the drug discovery process.

Untargeted analysis of NP extracts typically involves separation with liquid chromatography (LC) coupled with mass spectrometry (MS), which provides metabolic features, characterized by mass-to-charge (*m/z*) ratios measured in MS1 experiments, along with *m/z* values of molecular fragments (MS2 fragmentation)^3,4^ with a single metabolite typically detected as multiple metabolic features with the same retention time (RT) but different *m/z* values ^5,6^. A metabolite’s identity is determined by comparing its metabolite features with reference databases. However, manual annotation represents a daunting task, as it is time-consuming and dependent on expert knowledge, thus hampering scalability and high-throughput^7^. Subjectivity and human error may also introduce biases from researchers lacking expertise in particular classes of NPs. Addressing these limitations necessitates the use of standardized laboratory procedures, the application of automated annotation tools and appropriate reference databases.

The demand for high-throughput methods has led to the development of various tools to automatically annotate molecules present in complex mixtures. Here, we developed a comprehensive workflow, which integrates leading compound discovery tools (Compound Discoverer^TM^, MolDiscovery^9^ and MS2Query^10^), evaluates and ranks their prediction and integrates these with additional computational tools, biological criteria and metadata. We applied this pipeline to 7,400 extracts derived from 6,566 actinomycetes, leading to the annotation of 150,000 molecules, loaded in a web-based database called “Molecules Gateway”. The database, which can be employed for different searches, highlights the existence of a large, untapped chemical diversity from a relatively small number of actinomycetes.

## Results

### Design of the automated pipeline

We designed an automated annotation pipeline with the objective of performing systematic annotation of a large number of microbial fermentation extracts. After LC-HRMS analysis, the pipeline processes data through four sequential steps, as depicted in Figure 1A: 1) consolidation and filtering, whereby different *m/z* values originating from the same molecule are grouped into a single entity, followed by filtering out signals below fixed MS peak thresholds; 2) internal dereplication, whereby molecules identical to compounds observed in previously analyzed extracts do not enter the following steps, but are assigned the annotations of their matching molecules while the relevant data and metadata are added; 3) compound annotation, which exploits the annotation capabilities of Compound Discoverer, MS2Query and MolDiscovery, with their outputs harmonized and combined into a single output file; and 4) consistency and ranking, which weighs in the molecular formula (MF) predicted by SIRIUS^11^ and the RT calculated by jp^2^rt (https://github.com/mapio/jp2rt/tree/v0.2.3) and the biological consistency of the annotation (i.e., whether the molecule has been reported from the same biological source, actinomycetes in our case). At the end, experimental and calculated data, along with the associated metadata, enter a web-based, searchable database, the "Molecules Gateway", as described below.

**Figure 1:**
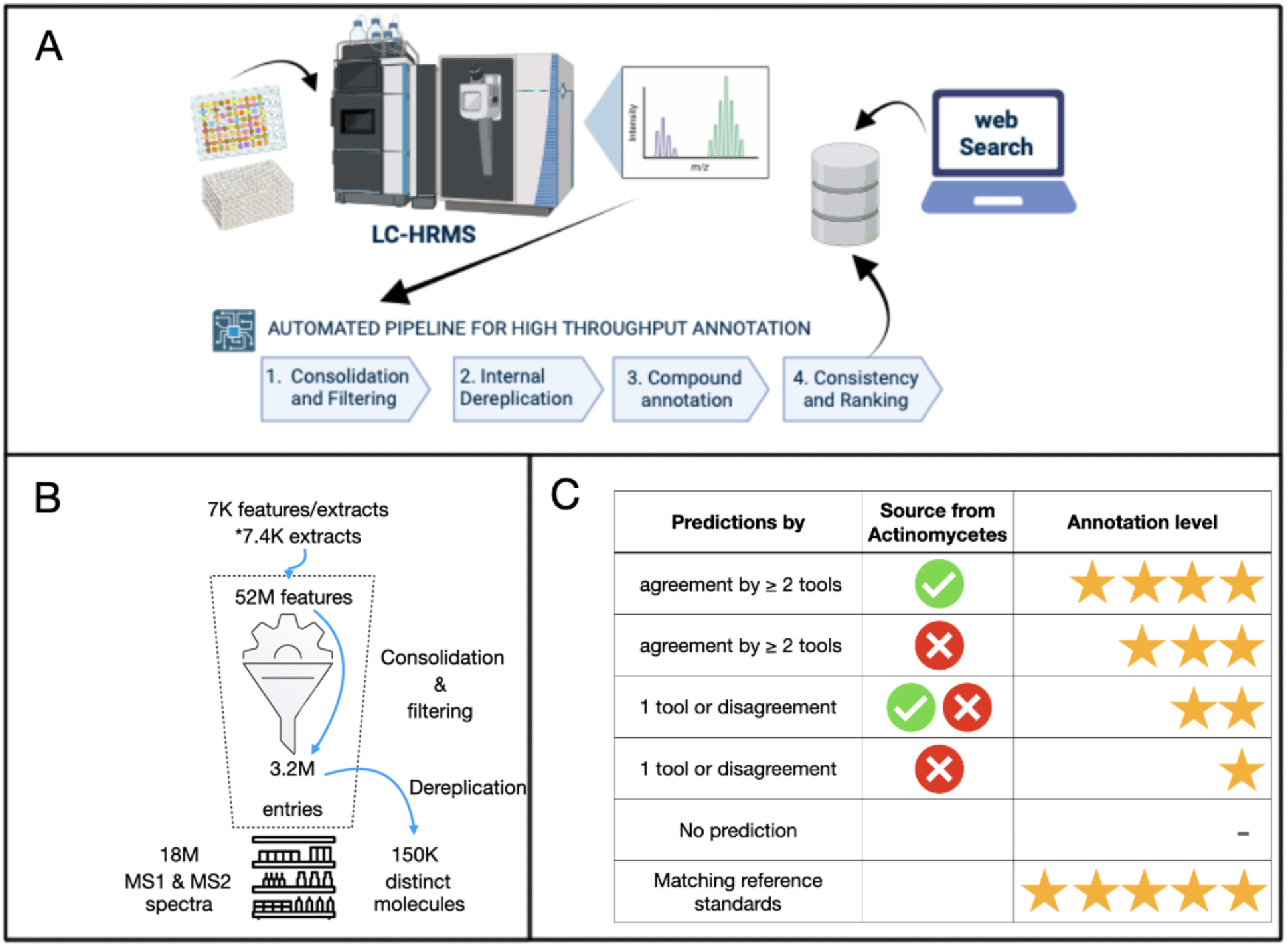
Panel A: Schematic of the process for analyzing microbial fermentation extracts and creating a web-searchable database, along with the logical flow of the annotation pipeline for processing HR-MS data. Panel B: summarized inputs and outputs after analyzing 7,400 actinomycete extracts. Entries denote unique molecule-extract combinations. Panel C: organization of annotation results into four different levels

### Concept validation

We evaluated the performance of Compound Discoverer, MolDiscovery and MS2Query in the annotation of a selected number of metabolites present in extracts derived from eight well-characterized *Streptomyces* strains: *S. coelicolor* and *S. avermitilis*, reference strains with extensive metabolite analyses^12^; *Streptomyces* sp. ID 38640^13^, *S*. *rimosus*, *S*. *mobaraensis*, *S*. *venezuelae*, *S. griseus* and *S*. *glaucescens* (see Table 1). Manual curation of the LC-HRMS data from the corresponding eight extracts led to the identification of 51 metabolites which were previously reported in the literature for these strains. In addition, we identified 8 molecules derived from the unfermented medium (see below). Overall, 22 molecules appeared in multiple extracts. When the HRMS raw data (on average, 7,000 features per extract) were processed through the first two steps of the annotation pipeline, different adducts originating from the same molecule (a total of 196 adducts from the 59 molecules) were correctly merged (Supplementary Table 1) and the 22 molecules appearing in multiple extracts were correctly dereplicated (Supplementary Table 2). Of note, the 59 molecules were always detected, at least in their most abundant *m/z* values, above a 10E7 peak area and generally showed a peak quality around 8. Next, we tested the performance of Compound Discoverer, MolDiscovery and MS2Query in annotating the 59 molecules. MolDiscovery exhibited the highest accuracy at 59%, followed by Compound Discoverer and MS2Query. Of note, each tool led to a significant number of false positives. The annotation outputs are reported in Supplementary Table 1 and summarized in Table 1. While no single tool was able to accurately identify all the 59 molecules, 46 of them (78%) were successfully identified by at least one tool, suggesting that an acceptable accuracy level could be achieved by a judicious choice of the most likely annotation.

**Table 1:**
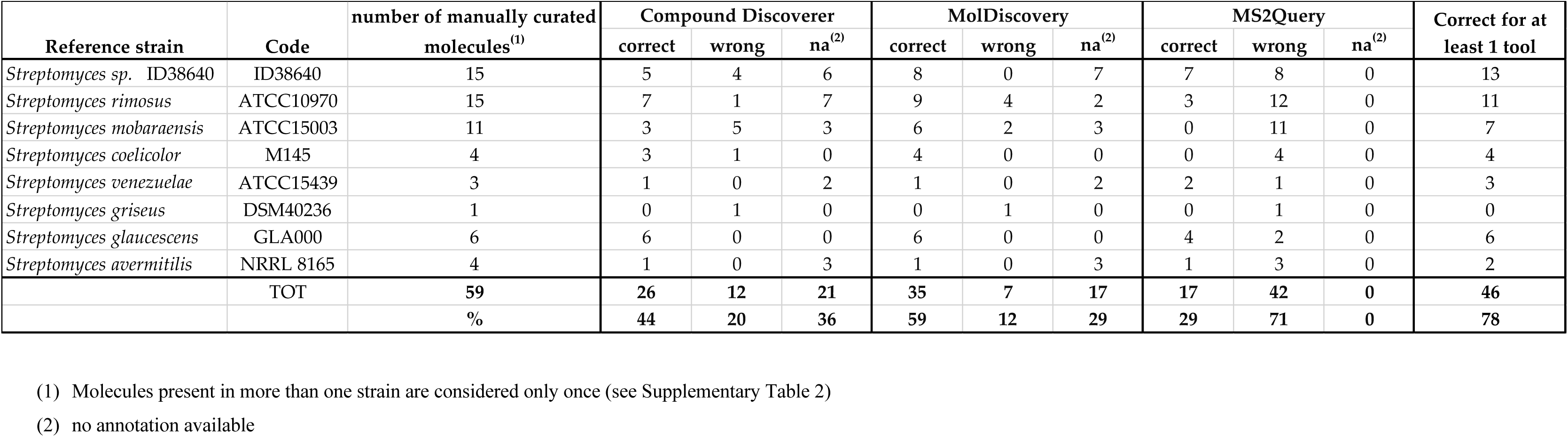
Performance of the annotation tools on validated molecules from reference strains. Consolidation and dereplication were performed by Compound Discoverer, and for this validation Compound Discoverer was limited to utilizing only the default databases (Chemspider, NPAtlas and mzCloud library). Signals corresponding to known molecules from the 8 reference strains were manually searched for within the three software outputs thus selecting MS1 *m/z* values and MS2 fragmentation scans related to the target molecules. For each software the correct and wrong prediction are reported together with the number of absence of prediction. The annotation was considered correct when it identified the exact molecule, an isomeric congener or, in the case of MS2Query, a highly similar molecule. The details for each molecule are listed in Supplementary Table 1. In the last column the number of correct annotations from at least one tool is reported for each reference strain. Molecules present in more than one strain are here considered only once but detailed in Supplementary Table 2.

### Implementation of the automated annotation pipeline

The annotation pipeline was implemented as a series of consecutive rounds of analysis and data processing, with each round consisting of 80 different extracts (i.e., the content of one 96-well microtiter plate suitable for bioassay screening). The input for each round of annotation was the raw HRMS data files, which underwent signal consolidation and filtering. By applying peak area and quality thresholds (see above) and by considering only signals with MS2 fragmentation, we observed a 10-fold reduction in the number of molecules entering step 2 (Figure 1A), with a great reduction in computational time, along with a lower risk of including MS artifacts. For dereplication, MS2 fragmentation data from newly analyzed extracts were compared with those present in the Dereplication Library, which initially included data from 452 reference molecules (Supplementary Table 3) and was augmented with all newly observed molecules at the end of each 80-extract round (see Methods). Molecules not matching entries in the Dereplication Library entered step 3 of the annotation pipeline, which involved parallel computational analysis utilizing Compound Discoverer, MolDiscovery, and MS2Query (see Methods for details). To manage the different output formats of the three tools, and compare and score their results into an automatic workflow, the Soupy software was developed (Supplementary Figure 1 and Supplementary Tables 4-5). In principle, each tool can annotate a molecule or not, thus from zero to three different structural predictions can be present. Instances without any prediction were labelled “unknown”, while molecules with one to three predictions entered the phase of “Consistency and Ranking”, which ultimately selected the most likely annotation. The criteria for consistency took into consideration the biological source, the SIRIUS-calculated MF and the jp^2^rt-calculated RT of the annotated molecules (Supplementary Table 6). Following a decision tree (Supplementary Figure 3), annotations were grouped into 12 different bins (Supplementary Table 6).

In order to evaluate its scalability, we applied the annotation pipeline to extracts derived from 6,566 strains from our collection of about 45,000 actinomycetes^14^. These strains - classified by partial 16S rRNA gene sequence - represent at least 86 genera belonging to 28 distinct actinomycete families (Supplementary Figure 4). Among the processed set, *Streptomyces* strains account for 2,000 isolates, and other important contributors are strains belonging to the families *Micromonosporaceae*, *Streptosporangiaceae*, *Thermomonosporaceae*, *Nocardiaceae* and *Pseudonocardiaceae*. Most strains were cultivated in a single fermentation medium, leading to 7,400 extracts prepared from the same number of cultures. Step 1 of the automated annotation pipeline reduced the complexity of the HR-MS data for each extract from about 7,000 features to about 400 grouped metabolite features (called molecules hereafter). Iterative dereplication of each 80-extract set against the Dereplication Library ultimately led to 150,777 distinct molecules representing 3,197,300 unique molecule-extract pairs. The average extract thus contributed 432 molecules, of which 20 unique. Overall, we processed over 52 million distinct MS features leading to a database of 18 million MS1 and MS2 spectra (Figure 1B).

To assess the reliability of the annotations, we manually curated 100 randomly selected molecules, specifically 20 for each set of grouped annotation bins (Supplementary Table 6). Manual curation involved, besides checking the consistency of the MS2 fragmentation with the proposed structure, using additional data (MS adducts and UV consistency, presence of congeners in the same extract, Molecular networking; see Methods) that were not considered during automated annotation. Apart from four instances with insufficient data to assist manual curation, we could categorize the automated annotations of the 96 molecules as likely or unlikely, as summarized in Table 2 (details in Supplementary Table 7). As expected, reliability was highest (75–85%) when two annotation tools agreed and the annotation was consistent with the biological source. Consistency with biological source was a more reliable indicator of annotation accuracy also when a single tool had annotated a molecule. On the basis of logical considerations and of the results from manual curation, the annotations were grouped into four levels that qualitatively provide an indication of their reliability (Figure 1C). When at least two tools agreed, annotations were scored with 4 stars when the biological source was coherent, and with 3 stars when not. When tools were in disagreement, or there was a prediction by a single tool, the annotation received two stars when coherent with the biological source or when at least two consistency were present, and one star when not. One-star annotation molecules were actually labelled as “undecided”, given the limited reliability of this category. Of note, among the 100 molecules, we observed three cases when possible duplicate molecules escaped the dereplication process (Supplementary Table 7). If this figure translates to a larger scale, a failure rate of 3% in dereplication would be quite acceptable.

**Table 2:** Results of the manual curation of 100 randomly selected molecules: 20 for each grouped annotation bins but 5a/b (undecided, as defined in Supplementary Table 6 and Supplementary Figure 2) and corresponding annotation levels. Deatails are reported in Supplementary Table 3.

| Group | Annotation bins | Source from Actinomycetes | Manual curation result |  |  | Annotation level (stars) |
| --- | --- | --- | --- | --- | --- | --- |
|  |  |  | Likely | Insufficient | Unlike |  |
| A | 1a: $\geq 2$ tools agreement (InChIKey) | yes | 17 (85%) | | 3 (15%) | 4 |
| C | 2a: $\geq 2$ tools agreement (similarity name) | | 15 (75%) | 2 (10%) | 3 (15%) | |
| B | 1b: $\geq 2$ tools agreement (InChIKey) | no | 5 (25%) | 2 (10%) | 13 (65%) | 3 |
| D | 2b: $\geq 2$ tools agreement (similarity name) | | 8 (40%) | | 12 (60%) | |
| E | 3a-c and 4a-c: 1 tool or disagreement | yes | 3 (15%) |  | 17 (85%) | 2 |

### The Molecules Gateway

The results from the automated annotation pipeline entered the Molecules Gateway, centered around the annotated molecules. Each molecule, identified by a specific ID, is accompanied by experimental data (detected adducts with their MS1 and MS2 data, associated area for each occurrence in different extracts, and RT) calculated data (see below) and associated metadata (a list of the extracts where it was identified, along with the code and genus of the extract- generating strain(s)), as shown in Figure 2. In addition, the Molecules Gateway flags whether a molecule matched any of the 1,031 distinct molecules detected in unfermented media (see Methods). For calculated data, the Molecules Gateway provides - for molecules classified at annotation levels 2 to 4, the proposed annotation, MF and chemical structure, InChIKeys, SMILES, name and synonyms, and NP classification. In addition, it includes the annotation calls made by each software, along with the consistency with the biological source, with the predicted versus experimental RT, and with the annotation-derived versus the SIRIUS-calculated MF. Molecules labelled as "undecided" contain all the information as above except for the proposed annotation, while molecules labelled as "unknown" include solely the *m/z* value, the experimental RT and, for masses below 850 Da, the SIRIUS-calculated MF.

**Figure 2:**
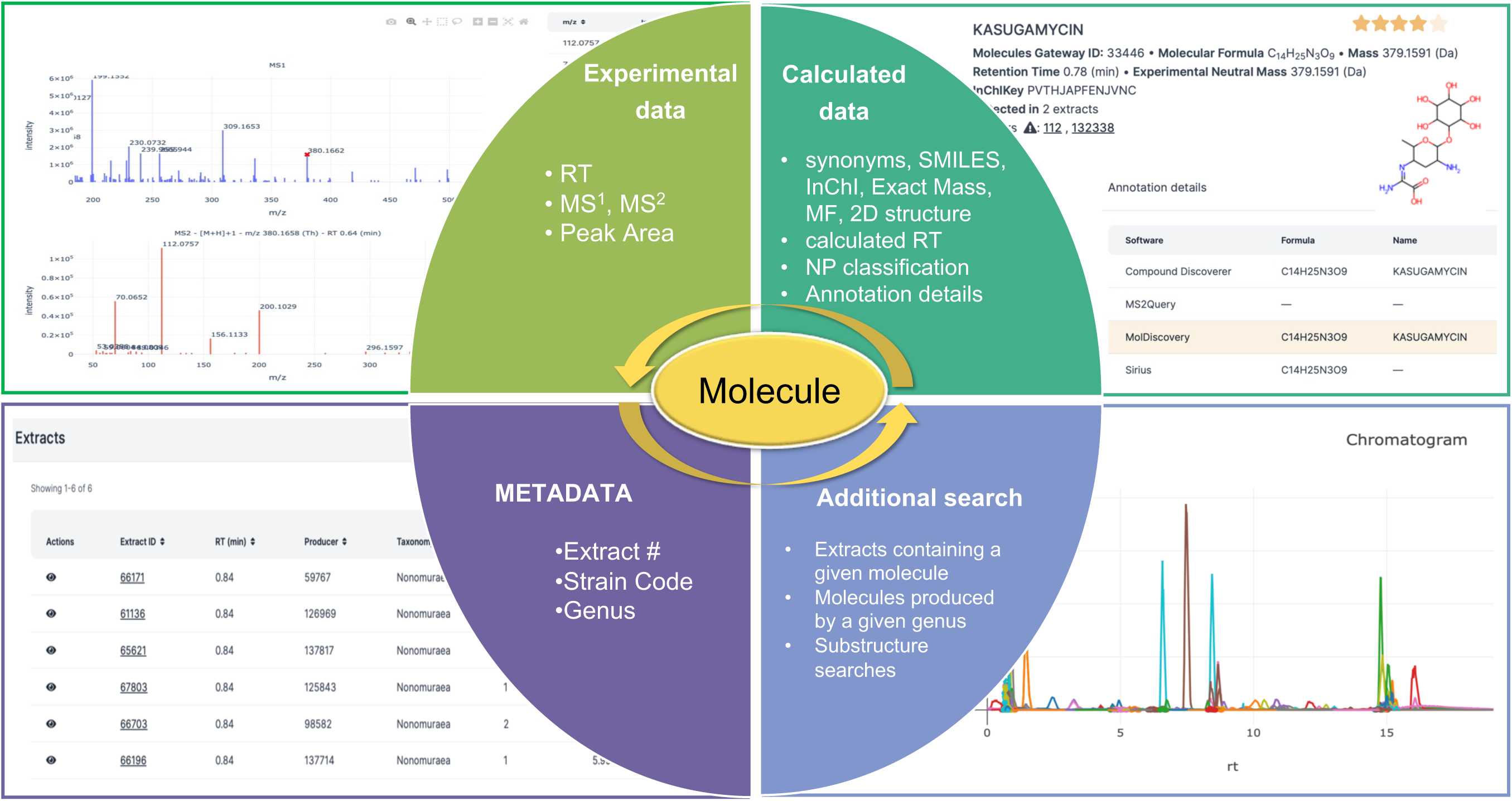
Schematic representation of the Molecules Gateway. All information linked to each molecule is interconnected. It includes experimental data (top left), calculated data (top right), metadata (bottom left) and additional search inputs and outputs (bottom right). A visual example is shown for each quadrant.

In addition, the Molecules Gateway provides a visual representation of the chromatogram for each extract, along with a list of all annotated molecules detected in each extract. This information can be useful for assessing the reliability of the annotation (e.g., by observing congeners of the annotated molecule) or for selecting the best strain for further work. Most experimental and calculated data are searchable in the Molecules Gateway, including substructure searches and MS2 fragment searches. Searches with experimental data span both known and unknown molecules, while searches with calculated data are limited to fully annotated molecules, with the exception of MFs, which can be performed for any molecule with exact mass below 850 Da.

### Using the Molecules Gateway

With its web-based, multiple search options, the Molecules Gateway can be used in many ways, ultimately leading to the identification of desired molecule(s), of the associated extract(s) and of the producing strain(s). We report below selected examples of using the Molecules Gateway, as schematized in Figure 3 and detailed in Supporting Data.

**Figure 3:**
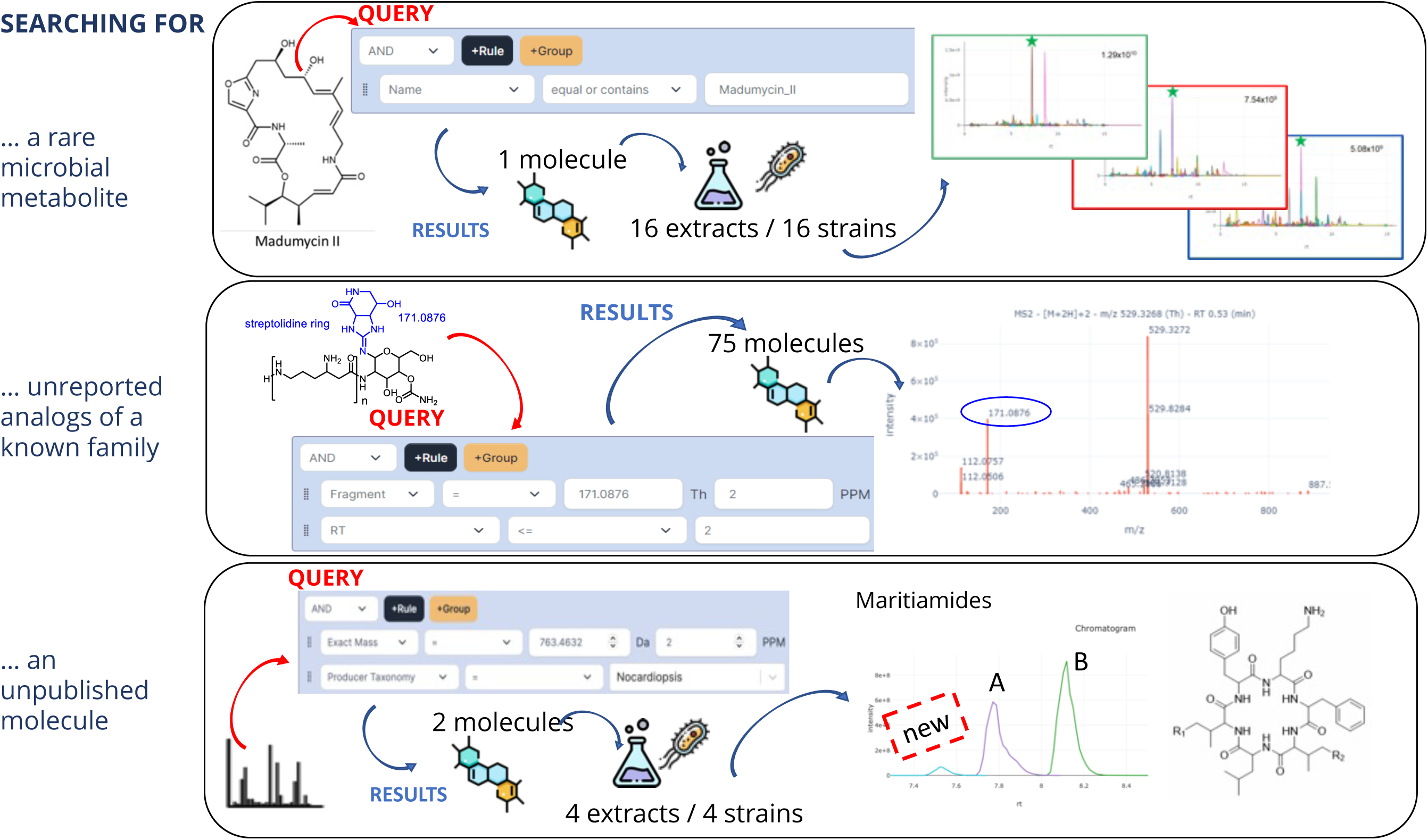
Possible uses of the Molecules Gateway. The figure shows the question asked, the search logics and an example of the outputs. Details for each example can be found as Supplementary Information as videos.

A first example involves searching for a rare microbial product, i.e., one with few commercial suppliers and few reports in the literature. Madumycin II, the simplest type A streptogramin, originally isolated from an *Actinoplanes*^15^ and a peptidyl transferase center inhibitor^16^, can fit into this category. Searching for “madumycin II” yields one hit (ID 56, with the highest reliability level) that appears in 16 distinct extracts, all from *Actinoplanes* strains. Selecting a specific extract leads to a web page with the corresponding chromatogram, displaying all detected molecules. This enables assessing the relative abundance of madumycin II, the presence of related molecules and the overall complexity of the metabolomes across the different strains, aiding in the selection of the most suitable strain for further work.

A second example pertains to searching for a yet undescribed analogs of a known metabolite family. As a specific case we took streptothricin, a molecule described in 1942^17^ and the subject of numerous studies^18^. Hallmarks of streptothricins are the streptolidine ring, which results in a diagnostic MS^2^ fragment of *m/z* 171.0876, and a high hydrophilicity, leading to low RTs in reverse phase chromatography. Searching the Molecules Gateway for "Fragment = 171.0876” and "RT ≤ 2 minutes” led to 75 hits, mostly from *Streptomyces* strains. Many of these hits contain *m/z* 171.0876 as a major MS2 fragment, as streptothricins do. In addition to several known members of the streptothricin family, many hits are labeled as “unknown”, suggesting they represent variants not yet reported in the scientific literature, even for a frequently occurring molecule known for many decades.

A third example consists in looking for a specific molecule not yet described in the scientific literature, e.g., a molecule under investigation in a scientist’s laboratory. To illustrate this scenario, we searched for maritiamides A and B^19^, antibacterial cyclic hexapeptides containing a Val and Ile residue, respectively, isolated from a *Nocardiopsis maritima* strain and reported in the scientific literature in 2024, thus absent from the libraries used in the present work. Searching for exact masses of 763.4632 and 777.4789 Da and restricting the search to the genus *Nocardiopsis* led to two hits (ID 69072 “undecided” and ID 69653 “unknown”), present in four distinct extracts, two from *Nocardiopsis* strains and two from *Streptomyces* strains. The fragmentation patterns of these hits are consistent with the maritiamides A and B structures, respectively. In addition, all extracts contain a third molecule (ID 69724 “unknown”) whose exact mass and fragmentation pattern are consistent with an additional maritiamide congener in which both Ile residues in maritiamide B are replaced by Val (Supplementary Figure 5). A noteworthy characteristic of the complex in that it consists of comparable amounts of maritiamide A and B in the *Nocardiopsis* extracts, while maritiamide A is the predominant form in the *Streptomyces* extracts (Supporting Figure 6).

The above examples illustrate how the Molecules Gateway can be readily queried to identify desired metabolites, analogs or unknown molecules, leading in many cases to the identification of multiple producer strains, also from different actinomycete genera. Having additional producing strains may help, for example, in the identification of the biosynthetic gene cluster, in detecting the presence or absence of analogs, in finding better producer strains, or in increasing the chances of finding a strain amenable to genetic manipulation.

### Exploring the chemical diversity from 6,566 Actinomycetes

The Molecules Gateway represents a dataset of 150,777 molecules annotated by the same workflow, produced by a diversified set of actinomycetes and present in homogeneously prepared extracts, which were analyzed under identical conditions. Thus, the dataset is amenable to different analyses that can provide insights into the chemical diversity of actinomycetes, including the existence of genus-specific and frequently occurring molecules.

In terms of annotation statistics, Compound Discoverer, MolDiscovery and MS2Query predicted molecules at very different rates due to their different annotation logics and the size and pertinence of the employed databases. Indeed, annotations ranged from 99% of call for Compound Discoverer to just 12% for MolDiscovery (Figure 4A). As expected from the large number of molecules, most molecules belong to the 0-star category (i.e., unknown), with about 60% of them having a SIRIUS-calculated MF (Figure 4B).

**Figure 4:**
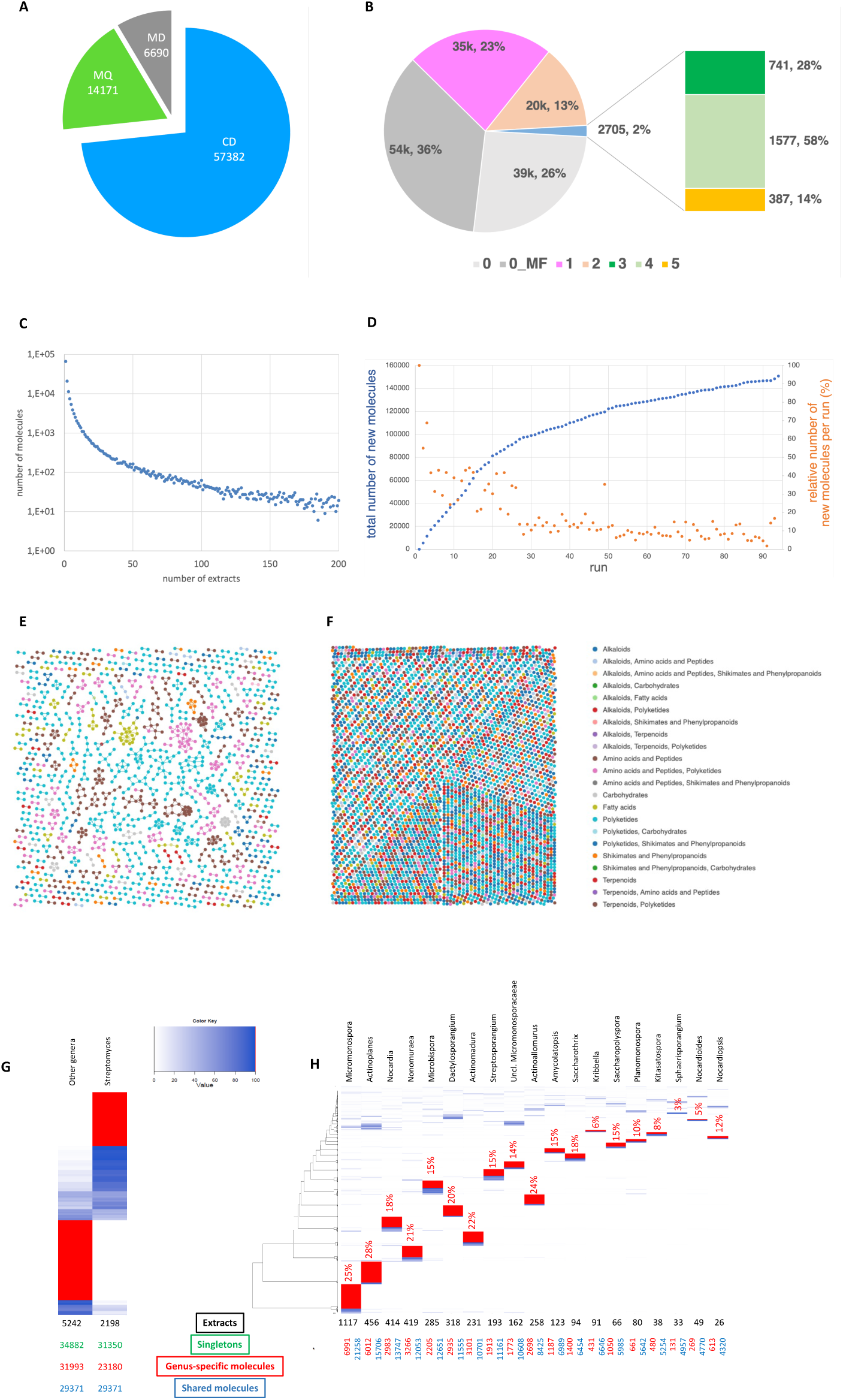
Overview of the 150,777 molecules listed in the Molecules Gateway. Panel A: Actual number of predictions by the different annotation tools. Panel B: Distribution of entries by annotation level. Panel C: Number of molecules found in a given number of extracts. Molecules present in >200 extracts are omitted. Panel D: Incremental growth of the Molecules Gateway. Panels E and F: chemical relatedness and originating biosynthetic class for 1,417 molecules arranged in families and 4,243 molecules as single nodes, respectively. Each dot indicates a molecule, and each color shows the originating biosynthetic class according to NPclassifier^42^. Clustering was performed using Morgan Fingerprint ^43^(with Radius set at 5) and Tanimoto Similarity^44^ (with cosine set at 0.7). Panels G and H: Heatmaps representing the distribution of molecules after UPGMA-based hierarchical clustering. Group-specific molecules are indicated by red bars, while gradient blue indicates increasing abundance for shared molecules. Panel G shows the distribution of molecules in *Streptomyces* and in the bulk of non-*Streptomyces* genera. The number of singletons is reported below in green type. Panel H shows the distribution of genus-specific molecules among the non-*Streptomyces* genera contributing at least 5,000 molecules. Panel H also reports the number of extracts (black type), the genus-specific molecules as number and percentage (red type), and the number of shared molecules (blue type) for each group. For both panels, annotated molecules, detected in more than then 2 and less than 500 extracts, were grouped according to genus and Ward-based hierarchical clustering in R (hclust). A distribution matrix, representing the specificity of each molecule to each genus, was used to create heatmaps in R-4.3.3 (R Core Team, 2023) using gplots and colorspace packages and a modified version of heatmap.3 function^45^ available at https://github.com/TV27/heatmap-metabo/blob/main/revheatmap3-R.

In terms of molecules distribution, just 3,120 out of 150K molecules in the Molecules Gateway were detected in more than 200 extracts and are likely to include, in addition to the 1,031 molecules present in the unfermented media, primary metabolites and frequently occurring specialized metabolites. The remaining molecules are distributed in the extracts in an exponentially decaying manner, as expected for specialized metabolites (Figure 4C). Looking at how the Dereplication Library increased in size after each 80-extract analysis can provide an indication of how further the database can grow by processing additional strains. During the first 20 runs, the number of new molecules increased at a rate of about 4,000 molecules per run, and then stabilized after the 30th run to about 1,200 per run, with no sign of a decreasing rate after 92 runs (Figure 4D). The relative number of new molecules added to the database varied with different runs, reflecting the varying contribution by different actinomycete genera. These data suggest that additional molecules can be added to the database at a rate of 10-20 new molecules per strain, depending on the chosen genus.

The distribution of exact masses and RTs for the 58,093 molecules with annotation shows that there is no obvious bias in RT and exact mass (Supplementary Figure 7). In terms of chemical diversity, Figures 4E and 4F report 1,417 molecules arranged in families and 4,243 molecules as single nodes, respectively, along with the corresponding NP classification. Overall, these data indicate that the annotation pipeline did not introduce any obvious bias, with molecular families or biosynthetic classes greatly over- or under-represented.

Finally, we wanted to establish the relative contribution of different genera to populating the Molecules Gateway. The number of contributed molecules varied greatly, from 199 molecules for *Embleya* (one single extract) to 83,901 molecules for *Streptomyces* (2,198 extracts), with a rough correlation between the number of extracts and the number of molecules (Figures 4G and 4H). We were also interested in metabolite distribution at genus level, which is the best taxonomic rank for comparative evaluation of biosynthetic potential^20^. Among the 83,901 molecules observed from *Streptomyces*, one third (29,371) are shared with other genera, while the rest are *Streptomyces*-specific (Figure 4G). When a similar analysis was performed on the remaining 96,246 molecules from non-*Streptomyces* genera, almost 70% of them (66,875) were not encountered in *Streptomyces*. Of note, the number of singletons (i.e., molecules observed in a single extract) was similar in both groups (Figure 4G). When the non-*Streptomyces* genera were analyzed individually, each genus contributed genus-specific metabolites ranging from 3% for *Sphaerisporangium* to 28% for *Actinoplanes* (Figure 4H). These data indicate that most if not all genera produce genus-specific molecules, while pointing to the most prolific genera in terms of genus-specific molecules. In this respect, genus diversification was empirically employed decades ago to reduce rediscovery rates in bioassay-based screens^21^.

## Discussion

After preparing 7,400 extracts derived from 6,566 actinomycetes and processing 52 million MS data, the Molecules Gateway with its 150,777 molecules represents one of the largest resources for analyzing and searching microbial products. The homogeneity of wet-lab and computational methods used to create the Molecules Gateway make it an attractive resource to accelerate research projects. Other databases of experimental MS data exist along with community-curated annotations: they range from 16- to 45-thousand molecules and include between 0.1 and 2 million MS data^22,23,24^. For example, Metabolights, GNPS/MassIVE and microbeMASST are valuable resources for the scientific community but their data remain intrinsically heterogeneous notwithstanding recent efforts at harmonization^25^. In addition, being community-based, their content reflects the interest of the scientific community and its willingness to contribute data, leading to possible biases. We believe the Molecule Gateway can nicely complement these resources, while providing a single place to access the physical samples that originated the data, along with the producing microorganisms.

Any MS-based annotation system suffers from the intrinsic limitation that stereoisomers and positional isomers cannot be distinguished, since they possess the same exact mass (MS1) and may display extremely similar or even undistinguishable mass fragmentations (MS2). Thus, in the absence of reference standards, stereoisomers and positional isomers will lead to identical annotations for molecules with (potentially) different RTs. This limitation is obviously present in the Molecules Gateway, notwithstanding our efforts at using reference standards. At the same time, the complexity of the automated annotation pipeline and the mere scale of the processed data, can lead to mis- annotations. For example, the utilized annotation tools have intrinsic limitations. Compound Discoverer operates on the largest database, which can be expanded but not tailor-reduced, resulting in many annotations that are not biologically relevant and unlikely to be correct. The accuracy of MS2Query, which operates on libraries of MS2 data from properly annotated molecules, is expected to increase when run against expanded and biologically relevant libraries. We observed that MolDiscovery is more reliable with some metabolite classes, e.g., unmodified peptides or oligosaccharides, that obey well-defined fragmentation rules. In addition, the annotation tools are known to suffer from an annotation bias that favors the known chemical space over the unknown^26^. Nonetheless, this annotation bias did not seem to have a profound effect on the annotations, as most molecules in the database belong to the unknown category (Fig. 4B). Finally, we observed that Compound Discoverer occasionally failed to properly dereplicate molecules as the size of the Dereplication Library increased, leading to redundant entries in the Molecules Gateway. Our sampling of 100 molecules suggests this phenomenon is limited to a small percentage. Notwithstanding these limitations, we believe the Molecules Gateway will be a useful tool, as experienced researchers can view experimental data and thus assess the validity of a suggested annotation and identify duplicated entries. Less experienced researchers can be guided by the annotation categories and by the consistency warnings associated with each molecule.

The Molecules Gateway contains far more molecules than those reported from all types of microorganisms after about 80 years of worldwide efforts^27^. The fact that most molecules are listed as unknown is thus not surprising, and literature data suggests that, on average, only around 10% of molecules can be annotated, especially in non-model organisms^26^. The accumulation curve we observe in Figure 4D is consistent with literature trends, which show an appreciable number of novel NPs discovered every year^28^. These data therefore suggest that thousands of additional molecules can be identified by processing the remaining portion of NAICONS’ library of actinomycetes or exploring additional fermentation media on the current strains. Genomic analyses suggest that only 3% of bacterial biosynthetic potential has been described, with *Streptomyces* as the most prolific actinomycete genus for potential undescribed metabolites^20^. As many genera are underrepresented in genomic databases, their metabolic potential could not be properly analyzed. Our metabolomic analysis suggest that some genera, for example *Actinoplanes*, *Micromonospora* and *Actinoallomurus* (Figure 4H), contribute a significant percentage of genus-specific molecules and can thus be considered prolific producers of potential novel metabolites, consistent with previous reports^21,29,30^.

While the annotation pipeline was employed to analyze actinomycete-derived extracts, it can be applied to extracts derived from other bacteria, from eukaryotic microorganisms or from higher organisms after the proper reference libraries are implemented to assist annotation and to establish biological consistency. This broadening of scope would empower the Molecules Gateway to furnish a universally applicable framework, where data searches for NPs from different biological sources can be readily associated with the availability of the corresponding chemical samples and biological material for further research. Such a comprehensive expansion may engender a transformative paradigm in the realm of NPs discovery and exploitation, fostering interdisciplinary collaborations and catalyzing innovation across many sectors.

## Methods

### WET-LAB PROCEDURES

#### Strain cultivation and extract preparation

The reference strains *Streptomyces coelicolor* M145, *S.avermitilis* NRRL 8165, S. *griseus* DSM 40236, *S. venezuelae* ATCC 15439, *Streptomyces* sp. ID38640, *S. rimosus* ATCC 10970, *S. mobaraensis* ATCC 15003 and *S. glaucescens* GLA000 and strains from the NAICONS collection were cultured as described^13^. For extract preparation, strains were cultivated for 3 days (*Streptomyces*) or for 7 days (other genera) at 30°C and 200 rpm. Production media used were: SV2^31^, M8, G1/0 and INA5^32^, M8acid (M8 with pH adjusted to 5.6), AF/A^33^ and Mare2Br (g/L; MgSO4.7H_2_O 12.3, NaCl 11.7, soluble starch 10, glucose.H_2_O 5, casein hydrolysate 2, CaCl_2_.2H_2_O 1.1, yeast extract 1, meat extract 1, CaCO_3_ 1, KCl 0.75, KBr 0.05, MnCl_2_.4H_2_O 0.0079, CuSO_4_.5H_2_O 0.0064, ZnSO_4_.7H_2_O 0.0015, FeSO_4_.7H_2_O 0.001).

For extract preparation, 4 mL ethanol was added to a 2-mL sample of each culture in a 15-mL centrifuge tube. The tube was shaken for 1 h at 30°C and centrifuged at 4000 rpm for 8 minutes. Then, 125 µl from each supernatant was transferred into individual wells of 96-well microtiter plates (80 extracts per well). Extracts from the seven unfermented production media were also prepared.

#### LC-HRMS analysis

Samples (8 µl) were analyzed using a Vanquish UHPLC system (Thermo Fisher Scientific) with a YMC-Triart ODS column (3.0 × 100 mm, S-1.9 μm, 12 nm) coupled to an Orbitrap Exploris™ 120 High-Resolution Mass Spectrometer (HRMS, Thermo Scientific Scientific) with Untargeted Data-Dependent Acquisition (DDA) analysis. The mobile phase, which was delivered at 0.8 ml min−1 at 40°C, consisted of 0.1% formic acid in H_2_O (A), LCMS-grade acetonitrile (B) and LCMS-grade isopropyl alcohol (C). The runtime sample analysis was 23 minutes and the mass to charge (m/z) ratio (MS1scan) was measured in the range from 150 to 2000. Further details as reported in Vind et al. 2023^34^.

### LIBRARIES and DATABASES

#### Dereplication Library

We made use of the customizable function of the Compound Discoverer suite (mzVault) to build a library of annotated compounds, which includes fragmentation patterns, adduct types, RT, SMILES and InChIKey of each molecule. The Dereplication Library initially consisted of 452 reference molecules: 92 reference standards and metabolites characterized in previous works and 360 manually curated molecules (Supplementary Table 3). After each annotation run the Dereplication Library was enriched with newly observed molecules and updated from the Molecule Gateway (see below).

#### MS2Query Library

The MS2Query default library was the GNPS^35^ library as of December 15, 2022, containing 314k MS2 fragmentation corresponding to 24k unique molecules. MS2Query was then trained on the default library expanded with data from the 452 reference molecules. [This step took 24 hours using a single light node (processor Intel E5-2683V4 2.1 GHz, 4 core and 8 GB of RAM) on the HPC INDACO computing platform (www indaco.unimi)].

### Bacterial Database (Bacterial-DB)

The database to run MolDiscovery was built by retrieving entries from NPAtlas^27^, Antimarin^36^ and ABL^37^ leading to 37,549 molecules originating from bacterial sources. All MS2 fragmentations were predicted by processing the database using the default MolDiscovery parameters.

### Natural Products Database (NP-DB)

This database, which includes 164k unique molecules, was created by merging data from NPAtlas^27^, Antimarin^36^, Coconut^38^, Lotus^39^ and ABL^37^, and includes key molecular attributes (SMILES, InChIKey, MF, names, exact mass, literature references and biological source, with 15,614 Actinomycete-derived molecules selectively flagged). The predicted RT for each molecule was calculated using jp^2^rt.

### Extract Database (Extract-DB)

It contains numerical codes for each extract associated to data on the producer strain code, its genus classification, the cultivation procedure and the plate number. The DB is used to retrieve and load metadata into the Molecules Gateway as described below.

### ANNOTATION WORKFLOW

**Tools validation** was performed as described in Table 1, Supplementary Table 1 and 2. SIRIUS^11^ (version 4.9.15) was tested on 219 reference molecules, and accurately calculated MFs for 100, 85 and 72% of molecules with masses up to 400, 600 and 850 Da, respectively (Supplementary Table 8).

**Soupy Software** In order to manage the entire workflow, we developed an ad-hoc software (named Soupy) that integrates several modules to support MS data analysis and signal annotation. The Soupy software was written in Python and the available commands are listed in Supplementary Table 4. Each of the commands (written in *italic*) is detailed in the following paragraphs. The entire annotation pipeline was implemented as a Snakemake workflow^40,41^ which allowed scalability and speed due to parallelization of the process. The Snakemake workflow was executed on Naicons Dell R620 server (SFF 2x E5-2630Lv2 with 6 cores, 2 threads per core, and 128GB RAM), processing one plate at a time. A total of 93 plates were analyzed using the iterative process reported in Supplementary Table 4. A "ThermoRawFileReader," (http://compomics.github.io/projects/ThermoRawFileParser) was accessed through a docker wrapper (https://www.compomics.com/) to achieve the conversion from raw files to mzML files. To facilitate data extraction from the Compound Discoverer output (cdResult file), Soupy incorporates the PyEDS library (https://github.com/thermofisherlsms/pyeds), which enables programmatic access to cdResults files. Additionally, Soupy utilizes RDKit, a comprehensive collection of computer chemistry and machine learning tools written in both C++ and Python (https://www.rdkit.org/) to clean up the data, including the conversion from SMILES to InChIKey.

**The annotation pipeline** (Fig. 1) was run as follows:

1. <u>The Consolidation and Filtering Step</u>: each 80-extract plate was analyzed together with three water:MeOH 1:1 blanks with the untargeted metabolomics workflow from Compound Discoverer (Version 3.3 SP2; Supplementary Figure 2). Parameters applied to each node of the workflow are available as Supplementary Information. Consolidation was performed by the “Group Compounds” node, which receives inputs from the following nodes: “Detect Compounds”, “Align timeTimes” and “Select Spectra”) using the default parameters. Filtering was performed on the Compound Discover output (.cdResult file) using the Soupy software (see below), selectively extracting the signals with RT 0.5 to 15 minutes, peak area ≥10^7^ and peak rating ≥8 (a Compound Discoverer parameter indicating the peak quality). Signals present in blanks were discarded.
2. <u>Internal Dereplication Step</u>: this involved using the “Group Compounds” and “Search mzVault” nodes - with the latter receiving input form the former - of Compound Discoverer (Supplementary Figure 2) to recognize the same molecule when present in different extracts of the same run (intra-run dereplication) and of different runs (inter-run dereplication using the Dereplication Library). Compounds that matched the Dereplication Library were exported by the *cdresult2matches* command as a matches.json file.
3. <u>The Annotation Step</u>: this was performed by Compound Discoverer using the original raw HRMS data files, using the “Assign Compound Annotation” node with parameters detailed in Supplementary Figure 2, adding the NP-DB to the “Search Mass Lists” node and assigning priority for matches inside this database. The computation time ranged from 2 to 9 hours. For running MolDiscovery and Mass2Query, the original raw HRMS data files were converted into mzML files by using the Soupy *raw2mzml* command (Supplementary Table 4). MolDiscovery^9^ (2.6.0-beta version) was launched via the Soupy *mzml2moldiscovery* command (Supplementary Table 4) by setting both the product ion and the parent mass threshold equal to 0.01 and by using the pre-processed Bacterial-DB. Running MolDiscovery took around 1 hour on a Dell R620 server (SFF 2x E5-2630Lv2 and 128 GB of RAM). MS2Query^10^ (Version 0.3) analysis was performed on each single .mzML file by applying default parameters and positive ion mode, and run on the HPC INDACO by the Soupy commands *mzml2indaco*, *slurm2indaco* and *indaco2ms2query* (Supplementary Table 4). A single light node was used for each mzML file with processor Intel E5-2683V4 2.1 GHz, 4 core and 8 GB of RAM. Running MS2Query took around 2 hours. MS2Query was implemented to account for exact matches only (within 2 ppm mass difference between experimental and theorical neutral mass).

After each annotation run, the Compound Discoverer outputs are an open-format SQLite database (.cdResult), where each molecule can have information ranging from a complete annotation, to MF only or no match. The compounds were extracted by the *cdresults2compound* command as a compounds.json file that contained MS1 and MS2 data, RT, exact mass, as well as, when available, compound name and predicted MF. The compounds.json file is the input for the *compound2summary* command, which completed and harmonized the information for each compound-based annotation exported from Compound Discoverer. MolDiscovery produces a single .tsv file for the 80 extracts and MS2Query individual .csv files for each extract. Additionally, the Compound Discoverer output is structured on compound-level logic (i.e., after signal consolidation and assigning a unique index named Compound_ID), while MolDiscovery and MS2Query operate at the MS2-scan level, potentially resulting in different annotations for the different adducts originating from a single compound. The MolDiscovery .tsv file (significant_matches_plate.tsv, see Supplementary Table 4) was the input for the *moldiscovery2summary* command, which cleaned up and harmonized the scan-based annotations selecting those with a score higher than 50. Since MS2Query renames the scan numbers from the raw files, the *mzml2summary* command exported a list of “extract – scan number pairs” for all MS2 fragmentation found in the MS2Query 80 mzML files and saved a single mzml.tsv summary file. Starting from the mzml.tsv summary and the 80 csv files, the *ms2query2summary* command produced a single ms2query.tsv summary which aligns the MS2Query output to each “extract – scan number pairs” in the original mzML file and ultimately cleaned up and harmonized the annotations from MS2Query possessing a score higher than 0.63.

SIRIUS was run, by using the output from the *cdresult2compound* command (compounds.json file) as input: the *compounds2mgf* command extracted an mgf file containing MS signals for molecules with molecular mass lower than 850 Dalton, irrespective of their annotation, and selecting protonated single charge adduct types only. Then, the *mgf2sirius* command run SIRIUS with the configuration reported in Supplementary Table 8 and the output file formula_identification.tsv, with predicted MFs and associated scores, was converted into a sirius.tsv file by *sirius2summary*, which filtered results for scores greater than 0 and converted the MF according to the Hill system.

The four .tsv summary files were the input for the *compute_annotations* command, which returned annotation on the filtered and unmatched compounds. The annotations were selected after aligning the summaries to squash multiple predictions and running the decision tree with consistency checks and ranking, as described below. For aligning the summaries, the outputs from each software were aligned using a multi-level index system (MultiIndex) in which the first level included the Compound_ID, the second level the Extract Number and the third level the Compound Discoverer-scan number (progressively numbered during the MS data acquisition). In order to retrieve the most reliable annotations from MolDiscovery and MS2Query (which perform scan-based annotations), multiple data were squashed into a single representative element by relying on four sequential criteria: highest binned score, SIRIUS agreement, most frequent InChiKey, lowest Compound Discoverer scan number. For score binning, MS2Query scores (scoring range [0.3,1]) were split into 20 uniform bins, while MolDiscovery scores (scoring range [50,220]) were split into 3 bins for scores under 100 and into 17 bins for scores above 100. (Fayyad and Irani, 1993). The MFs computed by MS2Query or MolDiscovery were individually compared with those predicted by SIRIUS, and scored as: 1 = agreement; -1 = disagreement; 0 = no predicted formula or disagreement when exact mass > 600 Da. For the InChIKey frequency, we considered the scans yielding the same InChIKey from a specific Compound Discoverer_ID, excluding scans with missing prediction. Finally, the annotation associated with the lowest Compound Discoverer scan number was preferred. Squashing resulted in a simple index based on Compound Discoverer_ID. An example is reported in Supplementary Table 5.

4. <u>Consistency and ranking Step</u>: the three criteria considered, the biological source, the MF and the RT of the annotated molecules, were part of the decision tree (Supplementary Figure 3, Supplementary Table 6), executed by the *compute_annotations* command. and resulting into 12 annotation bins. Each annotation resulted in a 3-tuple of values within {−1,0,1}. The biological consistency was rated as *1* when the biological source field in the NP-DB was True; *0* if the InChIKey was not present in the NP-DB or no InChiKey was assigned; or *−1* if the biological source field in the NP-DB was False. Similarly, the RT consistency was rated as *1* if the predicted RT was within 1.4 minutes of the experimental RT; *−1* if the difference exceeded 1.4 min; and *0* if no InChIKey was proposed or if the proposed InChiKey was not present in the NP-DB. The proposed MF was scored as *1* in case of agreement with the SIRIUS prediction; *−1* in case of disagreement; and *0* if at least one tool did not provide a MF. The arrangement aided in their lexicographic sorting, where consistency 3-tuples were categorized into (not mutually exclusive) classes as follows: class **T** comprised tuples with at least two occurrences of *1*, in any position; class **A** is a subset of **T**, with tuples with a *1* in the first position (i.e., the biological source consistency is *1*) and at least one additional *1*; class **O** comprises tuples with solely a *1* in the first position (Supplementary Table 6). These classes were employed in subsequent steps, to ascertain consistency winners. To determine the “winning” annotation, we also differentiated scenarios when an InChIKey was predicted from those when no prediction was made.

<u>The InChIKey match winner</u>: if the number of predicted InChiKeys was greater than 1, and 2 or 3 of them are identical, the winner was selected according to the name assigned, in the order, by MolDiscovery, Compound Discoverer, and MS2Query. The resulting annotation were assigned to bins **1a** and **1b**, depending on whether there was consistency or not, respectively, with the biological source (Supplementary Figure 3 and Supplementary Table 6).

<u>The Name similarity winner</u>: if the number of InChIKeys was greater than 1 but they were different, the predicted compound names were pairwise compared. If the similarity exceeded 75%, both names were included; after all the comparisons the set can contain 2 or 3 names. [Note that since similarity is not transitive, a list containing 3 names does not necessarily mean that they are all similar in pairs.] To assign the molecule’s name, the winner was subsequently determined according to the name assigned, in order, by MolDiscovery, Compound Discoverer, and MS2Query. The resulting annotation were assigned to bins **2a** and **2b**, depending on whether there was consistency or not, respectively, with the biological source (Supplementary Figure 3 and Supplementary Table 6).

<u>The consistency winners</u>: when the number of InChiKeys was greater than 1, but they were not identical and the name similarity was lower than 75%, a set of four winners was established as detailed in Supplementary Table 6. Accordingly, the annotation bins **3a**-**b**, **4a-b** were generated, together with **5a** (no consistency, or a single consistency different from biological source).

<u>The single InChIKey case</u>: if the number of InChIKey was 1 (i.e. only one annotation was proposed), the annotations were assigned to bin **3c when in** consistency class **A**, in bin **4c** when in consistency class **O** or in bin **5b** when there was no consistency with the biological source.

### MOLECULES GATEWAY

The web application Molecules Gateway was developed in Python using the Django framework using the PostgreSQL engine with the RDKit cartridge extension to store and search molecular structures. The database structure was defined using Django ORM, the *django-rdkit* library provided an implementation of the custom fields and functions of RDKit. The web application defers asynchronous long tasks to a distributed queue implemented using the Celery library. Using the process described in the previous steps, the following information is provided for every compound: the tool (Compound Discoverer, MolDiscovery or MS2Query) whose *prediction* is used in the final annotation; the *annotation bin*, a set of *warnings* that highlight discordant annotations or consistencies. Moreover, a total of 1031 molecules were identified and labeled as “component of unfermented media” by matching between the signals originated from extracts of unfermented media and the Reference Library obtained from all analyzed 6,5k extracts in the Molecules Gateway

Users can browse the Molecules Gateway using two search pages, simple and advanced. User queries are performed on a materialized view, a PostgreSQL table that acts like a computed cache to pre-aggregate and assemble data for user visualization. This view is manually refreshed using custom-developed scripts, as it takes one-two hours to produce the result. The advanced search is built using React.js, *react-querybuilder* for composing the UI, and *react- ketcher* to draw molecular structures; the query is then serialized as JSON and sent to Django. The query is validated using JSON Schema and then is converted to Django ORM to actually perform the search in the database.

**Data upload:** The *catalog_export* command compiled a plate.JSON file containing all the information to be imported into the Molecules Gateway. It received as input: the matches.json file produced by *cdresult2matches*; the compounds.json produced by *cdresults2compound*; the excel file produced by *compute_annotations*; the sirius.tsv file produced by *sirius2summary*; the directory housing the IDmap file generated by *vaults_export*. Finally, *catalog_export* appended the compounds lacking annotation within the “compound.json” labeling as "unknown" and appending the SIRIUS-generated MF when exact mass was lower than 850 Da. Additionally, Soupy used the information from comnpound.json file, about the presence of compounds in extracts, to query the Extracts-DB and retrieve the information on producer strain and its taxonomy.

The *Export2catalogue* command use the plate.JSON file produced by *catalog_export* command and the corresponding Compound Discoverer Results file (cdResult file) as inputs. For molecules matched against the Dereplication Library it appended the list of extracts in which they are detected, while new entries were created for novel annotation and unknown molecules with the list of extracts in which they were detected. MS1 and MS2 fragmentations are extracted from cdResult file for all the adducts. To avoid uploading wrongly assigned adducts, the theoretical neutral mass is calculated and if differs from the experimental one more than 2 ppm the adduct is discarded. After the upload to Molecule Gateway is complete, integrity checks, image generation of chemical structure (using RDKit software), NPclassification (using the REST API provided by NP Classifier^42^), and synonym search (using Pubchem REST APIs) tasks are launched.

**Updating the Dereplication Library from Molecules Gateway**: Dereplication Library was updated after completing each annotation and uploading run, using the *catalogue2vault* command to allow the next Dereplication step. For each entry the mzVault file contained the experimental MS2 fragmentations, RTs together with name, SMILES, InChIKey, measured neutral mass, precursor mass and related adduct type. To allow the tracking of the next matches, the same commands also generated a map file (IDmap) that correlate the unique Molecules Gateway IDs to the Dereplication Library indexes.

### QUALITY CONTROL OF ANNOTATIONS

In order to establish the reliability of the annotations in the Molecules Gateway, we grouped the annotation bins into six logical categories and randomly picked 20 equally-spaced molecule IDs within each of the categories excluding the last (bins **5a** and **5b**; Supplementary Table 6). Manual curation was performed by evaluating the following criteria: consistency of the MS2 fragmentation with the proposed chemical structure (categorized as not-consistent, not- plausible, plausible and consistent, or insufficient data); consistency of the detected adducts with the proposed chemical structure; presence of annotated congeners in the same extract; presence of a consistent UV spectrum in the chromatogram; and molecular networking in GNPS. For the last item, we exported a single MS2 fragmentation per each adduct type for the 22,912 molecules with chemical structure - divided in roughly equal subgroups on the basis of RT - and run a GNPS analysis (cosine ≥ 0.7). When one of the 20 molecules clustered in a network, the similarity in chemical structure was considered consistent with the annotation. Details are http://gnps.ucsd.edu/ProteoSAFe/status.jsp?task=af0dac3761cb4a46bba18f019cd4abcc; http://gnps.ucsd.edu/ProteoSAFe/status.jsp?task=da256bb97e8c405da4be1e5b53256e47; and http://gnps.ucsd.edu/ProteoSAFe/status.jsp?task=67b0d8e68e704323ad4f690ef610b536. Manual validation of MS2 fragmentation was used as the key criterion for evaluating accuracy of the annotation, with the additional criteria serving to reinforce or challenge the MS2 fragmentation outcomes. Consequently, MS2 fragmentation deemed "plausible/consistent" or "not-plausible/not-consistent" led to an evaluation of the annotation being either "likely" or "unlikely", respectively (Supplementary Table 7). Fragmentation data of the 100 molecules submitted to manual curation are available as downloadable mgf file in Zenodo (link).

## Supporting information

MS2 fragmentation data for 100 molecules for QC

MS2 fragmentation data for 48 molecules for QC

## Acknowledgements

We thank Armin Nourifar, Dora Ferreira, Christelle Yammine and Marco Barrili for help with extracts preparation; Mattia Monga, Luca Spampinato, for suggestions and stimulating discussions; Thierry Neus, Alessio Alessi and Mattia Ducci for informatic support.

## Author contributions

S.D., S.I.M., M.Si., M.I., P.M., M.So. designed the research. M.Sa. with M.Si, M.I., S.I.M. and S.D. developed Soupy and implement the automatic annotation pipeline. N.C., M.A. and M.C. developed and implemented Molecules Gateway. P.M., M.So, M.Si., M.I., S.I.M., A.T., S.S., C.B., T.V., A.G. and S.D performed the wet lab experiments and/or performed data analysis. S.I.M., M.Si, M.I. and P.M. wrote the manuscript, in cooperation with all other authors. J.J.J.vdH was involved in prolific discussion and reviewing the manuscript.

## Additional information

**Supplementary information** The online version contains supplementary material available at … Correspondence and requests for materials should be addressed to Sonia I. Maffioli.

## Data availability

All data are available in the main text and Supplementary Information.

MS1 and MS2 fragmentation data of 100 molecules considered for Quality Control are available as supplementary files.

Molecular Network used for quality control is available on GNPS (http://gnps.ucsd.edu/ProteoSAFe/status.jsp?task=af0dac3761cb4a46bba18f019cd4abcc; http://gnps.ucsd.edu/ProteoSAFe/status.jsp?task=da256bb97e8c405da4be1e5b53256e47; http://gnps.ucsd.edu/ProteoSAFe/status.jsp?task=67b0d8e68e704323ad4f690ef610b536).

Videos illustrating the described uses of the Molecules Gateway are available through the following link: https://micro4all.com/video-tutorial/

Interactive version of Figure 4 Panel E and F are available at the following links: https://micro4all.com/moldiv/ and https://micro4all.com/moldiv-singletons/, respectively.

The version of the heatmap.3 function used for figure 4 Panels G and H is available at https://github.com/TV27/heatmap-metabo/blob/main/revheatmap3-R.

## Code availability

The software jp^2^rt to predict RT is available through GitHub (https://github.com/mapio/jp2rt/tree/v0.2.3) or upon request. Software Soupy is available upon request.

## Supplementary Figures

**Supplementary Figure 1:** The annotation workflow. Distinct color fillings delineate the logical steps as in Figure 1. Yellow colors identify software, while items with blue outline denote steps managed by Soupy (see Methods). CD-, MD- and MQ- InChIKeys (initial 14 digits only) represent the molecules as annotated by Compound Discoverer, MolDiscovery and MS2Query, respectively. The output of the three annotation software was harmonized by Soupy in order to have the following data: compound ID, annotation name, extract number, retention time in minutes. Additionally, information regarding the biological source and the predicted RT were retrieved from the NP-DB, along with canonical SMILE, InChIKey (first 14-digits) and molecular formula expressed according to the Hill system^46^.

**Supplementary Figure 2:** Compound Discoverer Untargeted Metabolomics workflow, with default nodes to performs retention time alignment, unknown compound detection, and compound grouping across all samples, to predict elemental compositions for all compounds, to hide chemical background (using Blank samples), to identify compounds using mzCloud (MS2 fragmentation) and ChemSpider (formula or exact mass). Parameters applied for each node and additional databases (NP-DB) and nodes (mzVault for the Dereplication Library) are described in Methods and Supplementary Information.

**Supplementary Figure 3:** Decision tree that compares and ranks the annotation by Compound Discoverer (CD), MolDiscovery (MD) and MS2Query (MQ). Annotations generated by at least two software (left part of the decision tree) were compared based on InChIKey identity or, when absent, on similarity of compound names. Further evaluation focused on the consistency of the biological source (BS). When name similarity fell below 75% additional assessments were performed using consistency with BS, with Molecular Formula (MF) and with retention time (RT). When a single prediction was available (right part of the decision tree), its consistency was also evaluated on the basis of BS, MF and RT. The colored boxes denote the different annotation bins. A description of the decision-making process is provided in Methods and the annotation bins are detailed in Supplementary Table 6.

**Supplementary Figure 4:** Taxonomic distribution of extract-generating strains. The inner circle shows family level distribution, with families including <50 strains are merged in a single group. See legend for family color codes. The outer circle lists genera with more than 30 strains. Numbers in parentheses indicate number of strains.

**Supplementary Figure 5**: Chemical structures of maritiamides A and B^19^, the blue arrow indicates the variation point for the new identified congener. In panels A and B the fragmentations patterns and the HR-MS2 spectra are reported for maritiamides A and B respectively. In panels C, highlighted in grey, the fragmentation patterns and HR-MS2 spectrum are reported for the new identified congener.

**Supplementary Figure 6:** Maritiamides complex distribution by *Nocardiopsis* (extracts 58897 and 49939) and *Streptomyces* (extracts 32006 and 32007). In these conditions extract 58897 (highlighted in grey) from strain 92517 *Nocardiopsis* showed the highest productivity with a peak intensity of 4.59E8, ten time more than the other three extracts.

**Supplementary Figure 7:** Scatter plot correlating Retention time and exact mass for the 58,093 molecules annotated at the 1–5 confidence levels.

## Supplementary Tables (see Excel file)

**Supplementary Table 1** provides comprehensive insights into concept validation. It furnishes pertinent details for each molecule, including its strain ID, along with taxonomic information, the extract ID, and the corresponding ID allocated within the Molecules Gateway. Additionally, the table denotes the molecule’s name and specifies whether it is labeled as common metabolites from the medium components. The Manual Curation section delineates various criteria: the "Ref. STD" column indicates the availability of a standard for the molecule; "MS1&MS2" and "UV" fields verify mass fragmentation and UV spectrum consistency with that reported in literature for the molecule, respectively; "BGC" refers to the biosynthetic gene cluster presence in the genome. Furthermore, the table encompasses data such as total area, peak rating (a Compound Discoverer definition for mass peak shape and quality), molecular formula, exact mass, and retention time (RT). It also lists all detected adduct types for each molecule. Subsequently, the table shows the annotations proposed by each tool. Compound Discoverer offers name-based annotations, whereas MolDiscovery and MS2Query provide both name and score. For MS2Query, a column featuring the mass difference between experimental data and the query spectrum aids in distinguishing exact matches from analogues. Lastly, the "overall correct" column denotes whether the molecule was successfully identified by at least one tool, marked by a "y" for affirmative cases.

**Supplementary Table 2:** Concept validation. The 22 molecules appearing in multiple extracts are reported.

**Supplementary Table 3:** Reference molecules initially composing the Dereplication Library and including: 67 reference standards, 23 metabolites previously characterized from our strain collection^47,48,49,50,51,30,13,52,53,54^ and 362 manually curated molecules. MSI annotation level is also reported^55,56^. For molecules detected as multiple adducts one entry/adduct is listed.

**Supplementary Table 4:** List of Soupy commands with their corresponding input file, operation, output file and runtime.

**Supplementary Table 5:** example of squashing applied to annotation from MS2Query: for the same compound (ID_255 assigned by Compound Discoverer) MS2Query propose multiple annotations (one for each scan, in the same or different extracts). The single MS2Query annotation is selected by the following order: the highest score (after binning), the agreement with the Sirius Molecular Formula, the highest frequency, and the lowest scan number. The first two criteria result in a tie (we have six predictions with the same binned score (between 0.895 and 0.93) none of them in agreement with Sirius. The MS2Query annotation for ID_255 is thus selected among the most frequent predicted InChIKey (the first three ones) and among them the first one is chosen for its lowest scan number (in italic).

**Supplementary Table 6:** Summary of the criteria applied to rank the annotations from Compound Discoverer, MolDiscovery and MS2Query as described in Methods and illustrated in Supplementary Figure 3. (BS: Biological Source). The resulting 12 annotation bins are reported together with the grouped bins created for the Quality Control, and the resulting Annotation levels.

**Supplementary Table 7:** details about quality control summarized in Table 2. For each molecule its ID in Molecules Gateway and GNPS network are reported followed by the exact mass, the experimental RT, MF, the annotation level and the selected annotation. Then the annotation proposed by each tool in terms of name and InChIKey. Consistency checks are also reported: predicted RT, biological source ("y" when molecules are reported in literature to be produced by Actinomycetes, "n" when only other source are reported in the literature), calc. MF ("y" when the selected annotation has a MF in agreement with that calculated with SIRIUS, "n" when in disagreement). Manual curation evaluation: MS2 fragmentation check (five levels were judged from consistent, plausible, insufficient data, not- plausible and not-consistent), Adducts consistency (plausible or not-plausible), MS2 Network ("y" when the molecules is connected to structurally similar molecules, "n" when the molecule is connected to structurally unrelated molecules or is present as a single node), Congeners ("y" there are congeners of the molecule annotated in the same extract, "n" the contrary), UV ("y" when a consistent UV spectrum is coeluting to the mass signal) Overall results: ranking as likely, unlikely and insufficient data as described in the text.

**Supplementary Table 8:** SIRIUS test run on 219 Reference Molecules: Sirius correctly calculated MFs for 100, 85 and 72% of molecules with masses up to 400, 600 and 850 Da, respectively. Test molecules only contained [M+H]+ and [M+Na]+ adducts. SIRIUS was run with the following configuration: IsotopeSettings.filter = true; FormulaSearchDB = ””; Timeout.secondsPerTree = 0; FormulaSettings.enforced = HCNO; Timeout.secondsPerInstance = 0; AdductSettings.detectable = ”[[M + H]+]”; UseHeuristic.mzToUseHeuristicOnly = 650; AlgorithmProfile = orbitrap; IsotopeMs2Settings = SCORE; MS2MassDeviation.allowedMassDeviation = 2.0 ppm; NumberOfCandidatesPerIon = 3; UseHeuristic.mzToUseHeuristic = 300; FormulaSettings.detectable =B,Cl,Br,S; NumberOfCandidates = 10; AdductSettings.fallback = ”[[M + H]+]”; RecomputeResults = false)

## References

1. Newman, D. J. & Cragg, G. M. Natural Products as Sources of New Drugs over the Nearly Four Decades from 01/1981 to 09/2019. J. Nat. Prod. 83, 770–803 (2020).

2. De Medeiros, L. S. et al. Discovering New Natural Products Using Metabolomics-Based Approaches. in Microbial Natural Products Chemistry (ed. Pacheco Fill, T.) vol. 1439 185–224 (Springer International Publishing, Cham, 2023).

3. Bauermeister, A., Mannochio-Russo, H., Costa-Lotufo, L. V., Jarmusch, A. K. & Dorrestein, P. C. Mass spectrometry-based metabolomics in microbiome investigations. Nat. Rev. Microbiol. 20, 143–160 (2022).

4. Beniddir, M. A. et al. Advances in decomposing complex metabolite mixtures using substructure- and network-based computational metabolomics approaches. Nat. Prod. Rep. 38, 1967–1993 (2021).

5. Mathieu, N. & Patti, G. Systems-Level Annotation of a Metabolomics Data Set Reduces 25,000 Features to Fewer than 1,000 Unique Metabolites. Anal. Chem. 89, 10397–10406 (2017).

6. Dunn, W. B. et al. Mass appeal: metabolite identification in mass spectrometry-focused untargeted metabolomics. Metabolomics 9, 44–66 (2013).

7. Russo, F., Ottosson, F., Van Der Hooft, J. J. J. & Ernst, M. Deep Learning Models for LC-MS Untargeted Metabolomics Data Analysis. in *From Computational Logic to Computational Biology* (eds. Cantone, D. & Pulvirenti, A.) vol. 14070 128–144 (Springer Nature Switzerland, Cham, 2024).

8. Bach, E., Schymanski, E. L. & Rousu, J. Joint structural annotation of small molecules using liquid chromatography retention order and tandem mass spectrometry data. *Nat*. Mach. Intell. 4, 1224–1237 (2022).

9. Cao, L. et al. MolDiscovery: learning mass spectrometry fragmentation of small molecules. Nat. Commun. 12, 3718 (2021).

10. De Jonge, N. F. et al. MS2Query: reliable and scalable MS2 mass spectra-based analogue search. Nat. Commun. 14, 1752 (2023).

11. Dührkop, K. et al. SIRIUS 4: a rapid tool for turning tandem mass spectra into metabolite structure information. Nat. Methods 16, 299–302 (2019).

12. Nett, M., Ikeda, H. & Moore, B. S. Genomic basis for natural product biosynthetic diversity in the actinomycetes. Nat. Prod. Rep. 26, 1362 (2009).

13. Iorio, M. et al. Blocks in the pseudouridimycin pathway unlock hidden metabolites in the Streptomyces producer strain. Sci. Rep. 11, 5827 (2021).

14. Monciardini, P., Iorio, M., Maffioli, S., Sosio, M. & Donadio, S. Discovering new bioactive molecules from microbial sources. Microb. Biotechnol. 7, 209–220 (2014).

15. Chamberlin, J. W. & Chen, S. A2315, new antibiotics produced by Actinoplanes philippinensis. 2. Structure of A2315A. *J. Antibiot. (Tokyo)* **30**, 197–201 (1977).

16. Osterman, I. A. et al. Madumycin II inhibits peptide bond formation by forcing the peptidyl transferase center into an inactive state. Nucleic Acids Res. 45, 7507–7514 (2017).

17. Waksman, S. A. & Woodruff, H. B. Streptothricin, a New Selective Bacteriostatic and Bactericidal Agent, Particularly Active Against Gram-Negative Bacteria. Exp. Biol. Med. 49, 207–210 (1942).

18. Franck, E. & Crofts, T. S. History of the streptothricin antibiotics and evidence for the neglect of the streptothricin resistome. Npj Antimicrob. Resist. 2, 3 (2024).

19. Lee, J., Um, S., Kim, E.-H. & Kim, S. H. Genomic and Metabolomic Analyses of *Nocardiopsis maritima* YSL2 as the Mycorrhizosphere Bacterium of *Suaeda maritima* (L.) Dumort. J. Nat. Prod. 87, 733–742 (2024).

20. Gavriilidou, A. et al. Compendium of specialized metabolite biosynthetic diversity encoded in bacterial genomes. Nat. Microbiol. 7, 726–735 (2022).

21. Parenti, F. & Coronelli, C. Members of the Genus Actinoplanes and their Antibiotics. Annu. Rev. Microbiol. 33, 389–411 (1979).

22. Yurekten, O. et al. MetaboLights: open data repository for metabolomics. Nucleic Acids Res. 52, D640–D646 (2024).

23. Leao, T. F. et al. Quick-start infrastructure for untargeted metabolomics analysis in GNPS. Nat. Metab. 3, 880– 882 (2021).

24. Zuffa, S. et al. microbeMASST: a taxonomically informed mass spectrometry search tool for microbial metabolomics data. Nat. Microbiol. 9, 336–345 (2024).

25. El Abiead, Y., et al. Enabling pan-repository reanalysis for big data science of public metabolomics data. Preprint at 10.26434/chemrxiv-2024-jt46s (2024).

26. De Jonge, N. F. et al. Good practices and recommendations for using and benchmarking computational metabolomics metabolite annotation tools. Metabolomics 18, 103 (2022).

27. van Santen, J. A. et al. The Natural Products Atlas 2.0: a database of microbially-derived natural products. Nucleic Acids Res. 50, D1317–D1323 (2022).

28. Pye, C. R., Bertin, M. J., Lokey, R. S., Gerwick, W. H. & Linington, R. G. Retrospective analysis of natural products provides insights for future discovery trends. Proc. Natl. Acad. Sci. 114, 5601–5606 (2017).

29. Wagman, G. H. Antibiotics from Micromonospora. *Annu. Rev. Microbiol.* 34, 537–558 (1980).

30. Iorio, M. et al. Allopeptimicins: unique antibacterial metabolites generated by hybrid PKS-NRPS, with original self-defense mechanism in *Actinoallomurus*. RSC Adv. 12, 16640–16655 (2022).

31. Zettler, J. et al. New Aminocoumarins from the Rare Actinomycete *Catenulispora acidiphila* DSM 44928: Identification, Structure Elucidation, and Heterologous Production. ChemBioChem 15, 612–621 (2014).

32. Donadio, S., Monciardini, P. & Sosio, M. Chapter 1 Approaches to Discovering Novel Antibacterial and Antifungal Agents. in *Methods in Enzymology* vol. 458 3–28 (Elsevier, 2009).

33. Cruz, J. C. S. et al. Brominated Variant of the Lantibiotic NAI-107 with Enhanced Antibacterial Potency. J. Nat. Prod. 78, 2642–2647 (2015).

34. Vind, K. et al. Megalochelin, a Tridecapeptide Siderophore from a Talented Streptomycete. ACS Chem. Biol. 18, 861–874 (2023).

35. Wang, M. et al. Sharing and community curation of mass spectrometry data with Global Natural Products Social Molecular Networking. Nat. Biotechnol. 34, 828–837 (2016).

36. Blunt, J., Munro, M. & Laatsch, H. AntiMarin database. (2006).

37. Simone, M., Monciardini, P., Gaspari, E., Donadio, S. & Maffioli, S. I. Isolation and characterization of NAI-802, a new lantibiotic produced by two different Actinoplanes strains. J. Antibiot. (Tokyo) 66, 73–78 (2013).

38. Sorokina, M., Merseburger, P., Rajan, K., Yirik, M. A. & Steinbeck, C. COCONUT online: Collection of Open Natural Products database. J. Cheminformatics 13, 2 (2021).

39. Rutz, A. et al. The LOTUS initiative for open knowledge management in natural products research. eLife 11, e70780 (2022).

40. Mölder, F. et al. Sustainable data analysis with Snakemake. F1000Research 10, 33 (2021).

41. Köster, J. & Rahmann, S. Snakemake—a scalable bioinformatics workflow engine. Bioinformatics 28, 2520–2522 (2012).

42. Kim, H. W. et al. NPClassifier: A Deep Neural Network-Based Structural Classification Tool for Natural Products *J*. Nat. Prod. 84, 2795–2807 (2021).

43. Morgan, H. L. The Generation of a Unique Machine Description for Chemical Structures-A Technique Developed at Chemical Abstracts Service. J. Chem. Doc. 5, 107–113 (1965).

44. Bajusz, D., Rácz, A. & Héberger, K. Why is Tanimoto index an appropriate choice for fingerprint-based similarity calculations? J. Cheminformatics 7, 20 (2015).

45. Griffith, D. M., Veech, J. A. & Marsh, C. J. **cooccur** : Probabilistic Species Co-Occurrence Analysis in *R*. J. Stat. Softw. 69, (2016).

46. Hill, E. A. On a system of indexing chemical literature; adopted by the classification division of the u. S. Patent office. ^1^. J. Am. Chem. Soc. **22**, 478–494 (1900).

47. Vind, K. et al. Megalochelin, a Tridecapeptide Siderophore from a Talented Streptomycete. ACS Chem. Biol. 18, 861–874 (2023).

48. Vind, K., et al. *N* -Acetyl-Cysteinylated Streptophenazines from *Streptomyces*. J. Nat. Prod. 85, 1239–1247 (2022).

49. Pozzi, R. et al. The genus Actinoallomurus and some of its metabolites. J. Antibiot. (Tokyo) 64, 133–139 (2011).

50. Mazzetti, C. et al. Halogenated Spirotetronates from *Actinoallomurus*. J. Nat. Prod. 75, 1044–1050 (2012).

51. Maffioli, S. I. et al. Antibacterial nucleoside-analog inhibitor of bacterial RNA polymerase: pseudouridimycin. Preprint at 10.1101/106906 (2017).

52. Iorio, M. et al. Novel Polyethers from Screening Actinoallomurus spp. Antibiotics 7, 47 (2018).

53. Iorio, M. et al. Antibacterial Paramagnetic Quinones from *Actinoallomurus*. J. Nat. Prod. 80, 819–827 (2017).

54. Iorio, M. et al. Chrolactomycins from the Actinomycete *Actinospica*. J. Nat. Prod. 75, 1991–1993 (2012).

55. Schymanski, E. L. et al. Identifying Small Molecules via High Resolution Mass Spectrometry: Communicating Confidence. Environ. Sci. Technol. 48, 2097–2098 (2014).

56. Sumner, L. W. et al. Proposed minimum reporting standards for chemical analysis: Chemical Analysis Working Group (CAWG) Metabolomics Standards Initiative (MSI). Metabolomics 3, 211–221 (2007).

