## Supplementary Figures for "The Molecules Gateway: a homogeneous, searchable database of 150k annotated molecules from Actinomycetes"

**Supplementary Figure 1:** The annotation workflow. Distinct color fillings delineate the logical steps as in Figure 1. Yellow colors identify software, while items with blue outline denote steps managed by Soupy (see Methods). CD-, MD- and MQ- InChIKeys (initial 14 digits only) represent the molecules as annotated by Compound Discoverer, MolDiscovery and MS2Query, respectively. The output of the three annotation software was harmonized by Soupy in order to have the following data: compound ID, annotation name, extract number, retention time in minutes. Additionally, information regarding the biological source and the predicted RT were retrieved from the NP-DB, along with canonical SMILE, InChIKey (first 14-digits) and molecular formula expressed according to the Hill system[46].

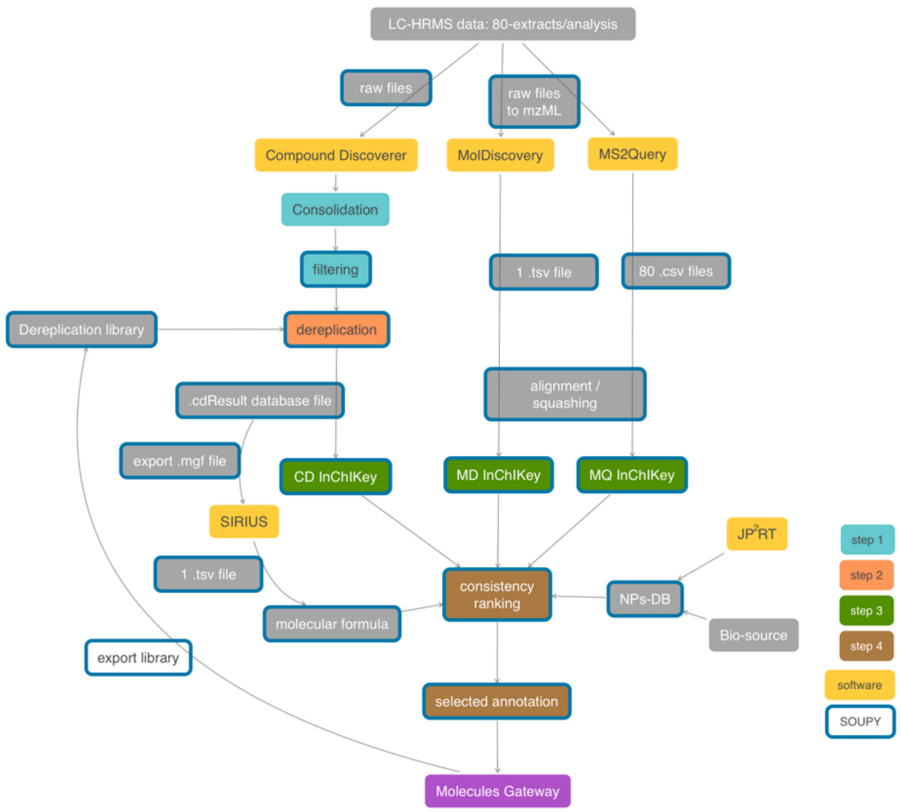

**Supplementary Figure 2:** Compound Discoverer Untargeted Metabolomics workflow, with default nodes to performs retention time alignment, unknown compound detection, and compound grouping across all samples, to predict elemental compositions for all compounds, to hide chemical background (using Blank samples), to identify compounds using mzCloud (MS2 fragmentation) and ChemSpider (formula or exact mass). Parameters applied for each node and additional databases (NP-DB) and nodes (mzVault for the Dereplication Library) are described in Methods and Supplementary Information.

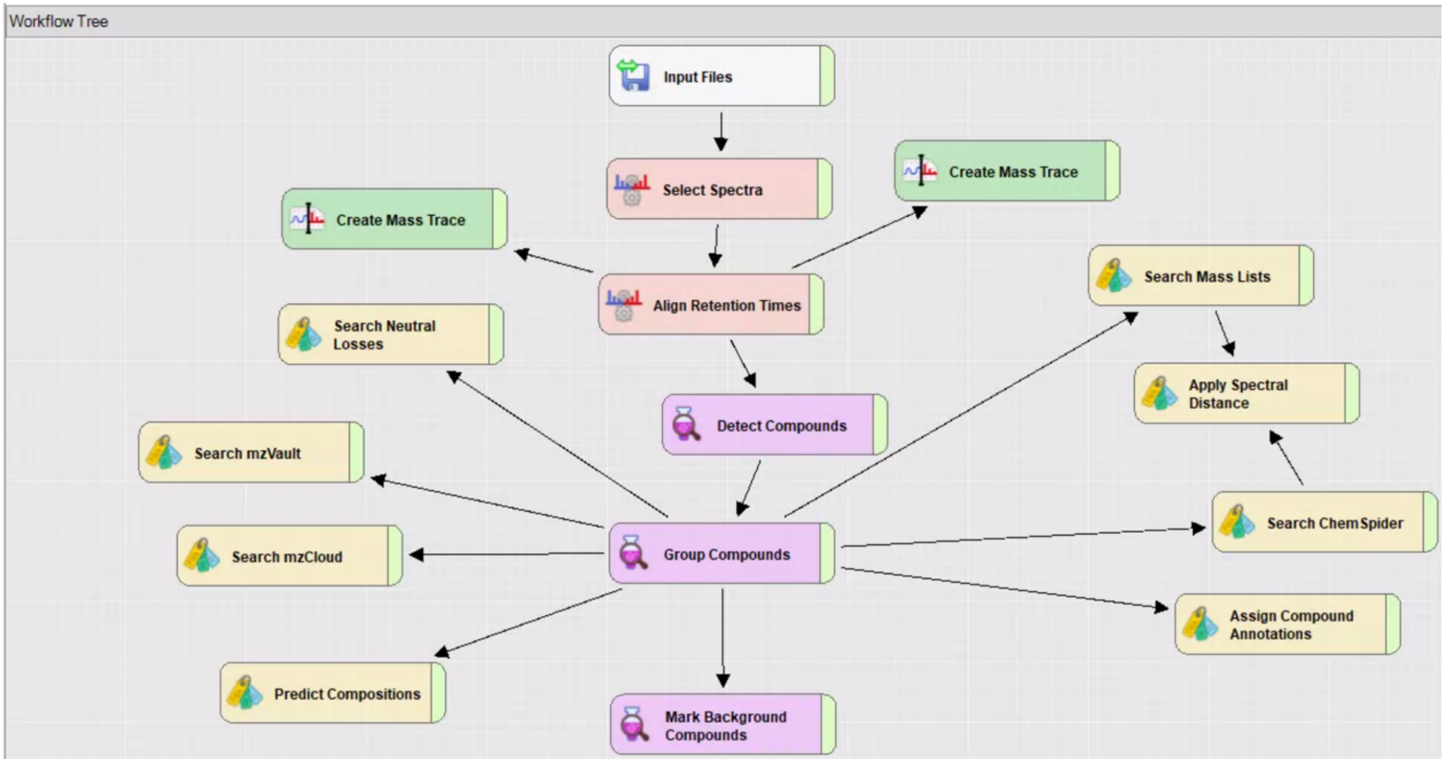

**Supplementary Figure 3:** Decision tree that compares and ranks the annotation by Compound Discoverer (CD), MolDiscovery (MD) and MS2Query (MQ). Annotations generated by at least two software (left part of the decision tree) were compared based on InChIKey identity or, when absent, on similarity of compound names. Further evaluation focused on the consistency of the biological source (BS). When name similarity fell below 75% additional assessments were performed using consistency with BS, with Molecular Formula (MF) and with retention time (RT). When a single prediction was available (right part of the decision tree), its consistency was also evaluated on the basis of BS, MF and RT. The colored boxes denote the different annotation bins. A description of the decision-making process is provided in Methods and the annotation bins are detailed in Supplementary Table 6.

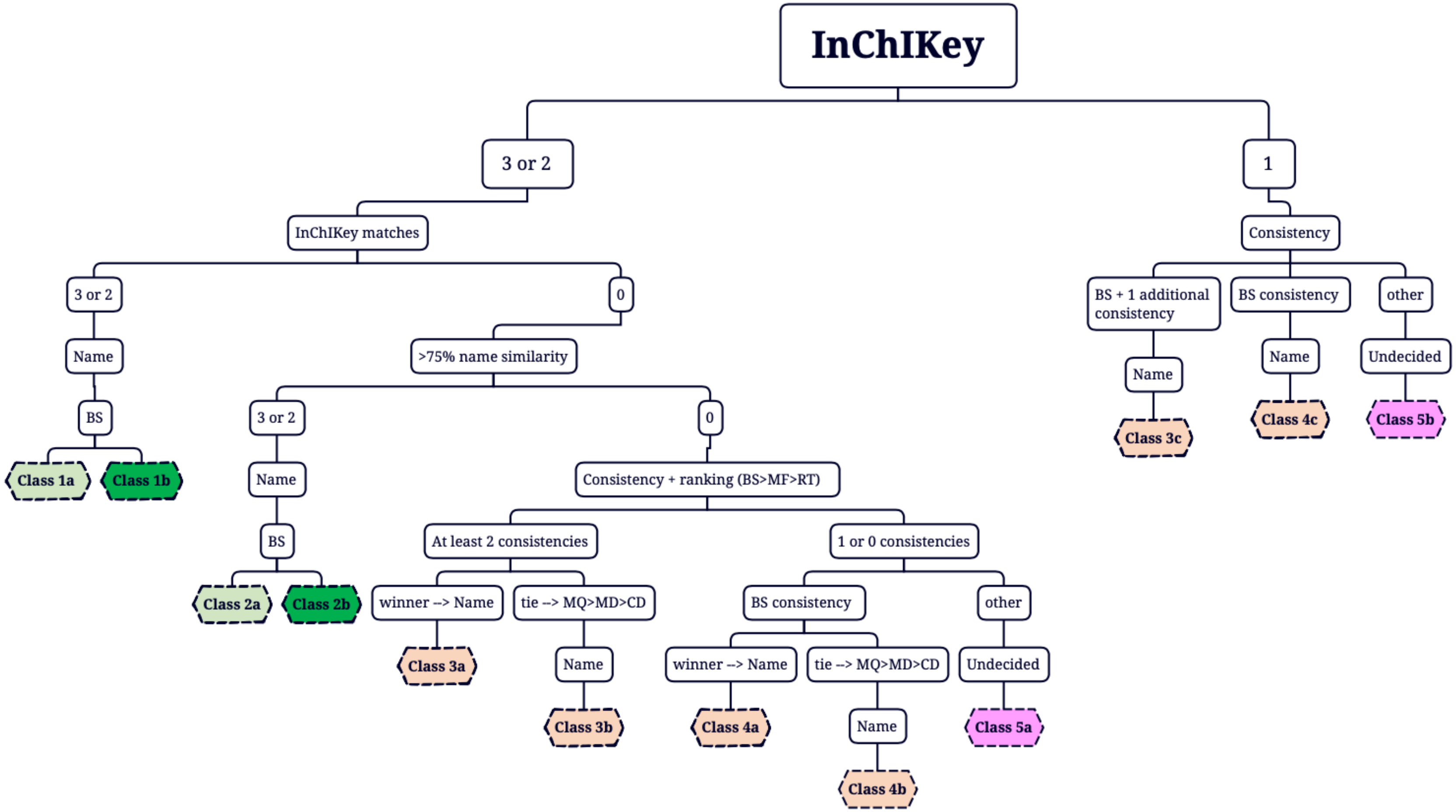

**Supplementary Figure 4:** Taxonomic distribution of extract-generating strains. The inner circle shows family level distribution, with families including <50 strains are merged in a single group. See legend for family color codes. The outer circle lists genera with more than 30 strains. Numbers in parentheses indicate number of strains.

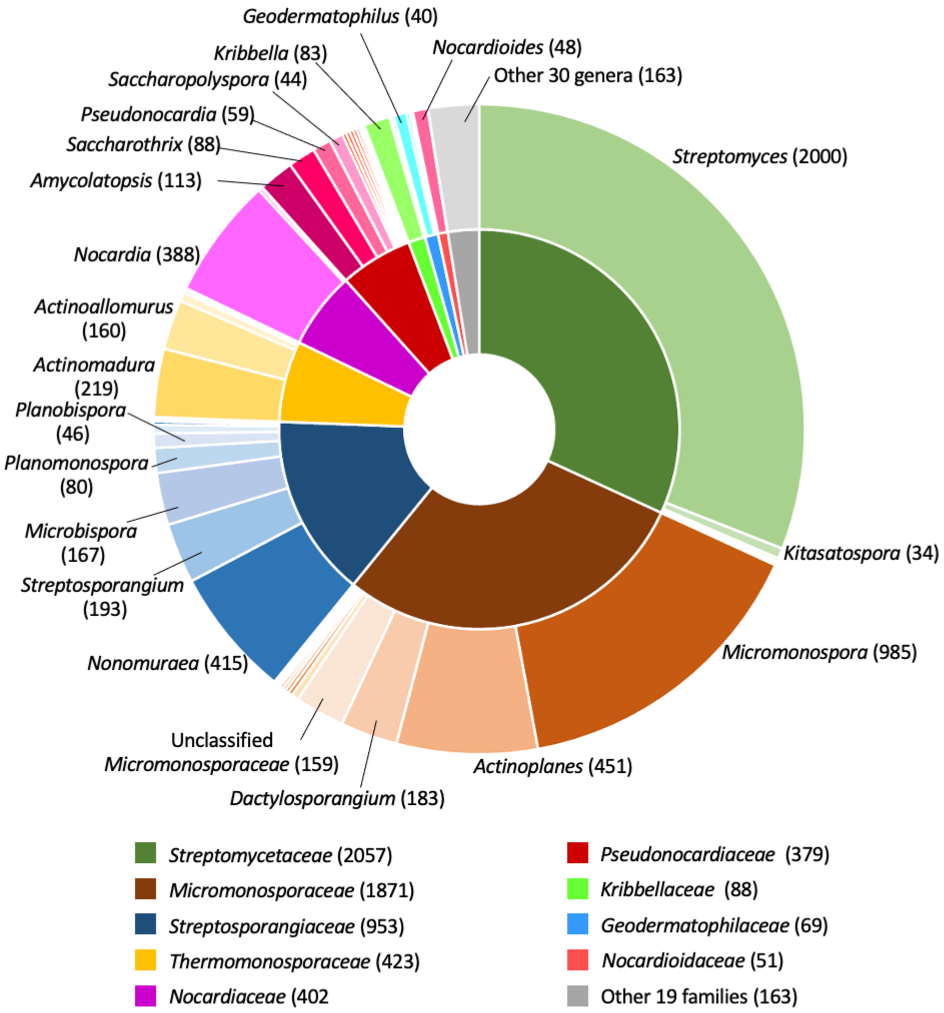

**Supplementary Figure 5:** Chemical structures of maritiamides A and B[19], the blue arrow indicates the variation point for the new identified congener. In panels A and B the fragmentations patterns and the HR-MS2 spectra are reported for maritiamides A and B respectively. In panels C, highlighted in grey, the fragmentation patterns and HR-MS2 spectrum are reported for the new identified congener.

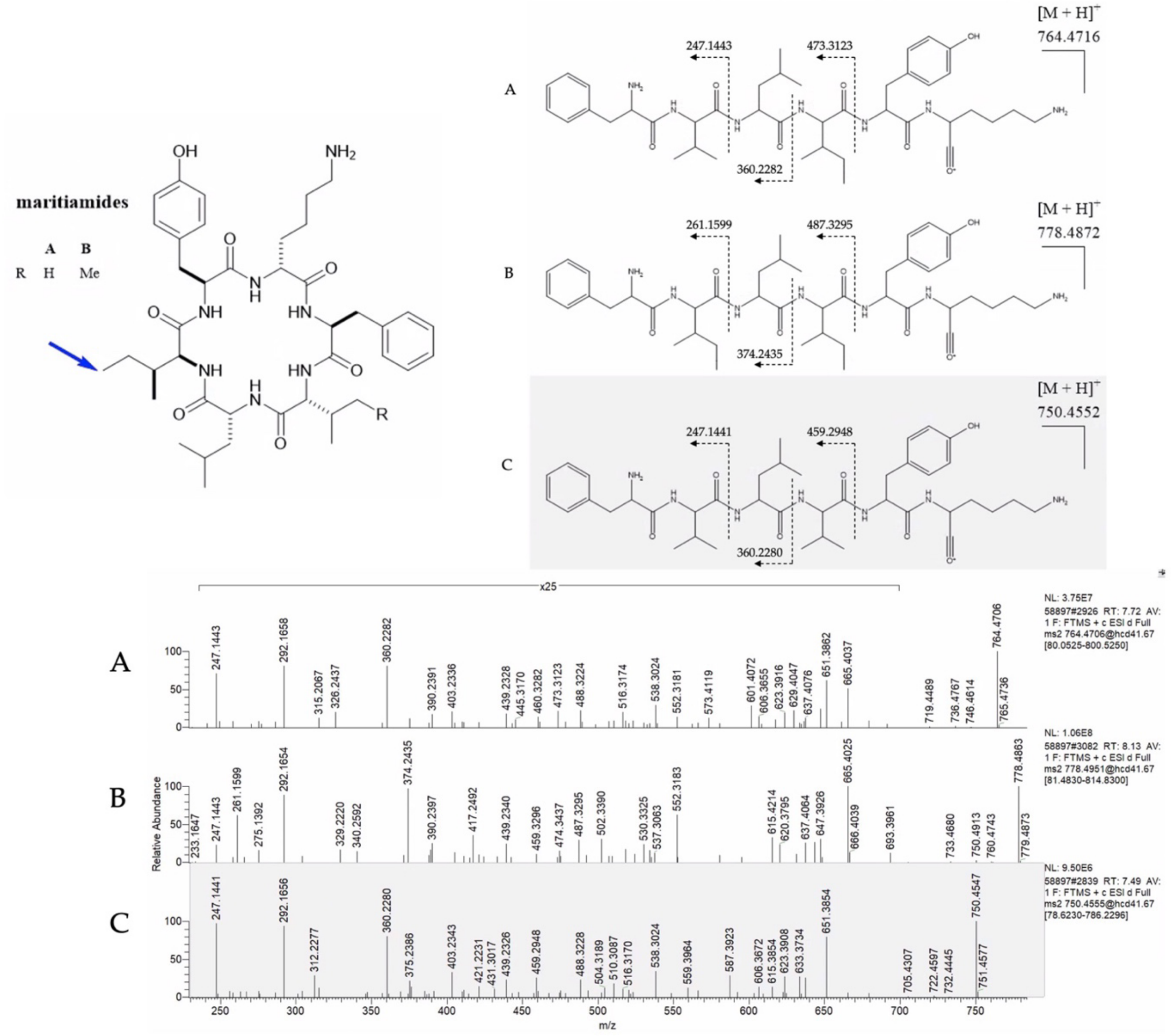

**Supplementary Figure 6:** Maritiamides complex distribution by *Nocardiopsis* (extracts 58897 and 49939) and *Streptomyces* (extracts 32006 and 32007). In these conditions extract 58897 (highlighted in grey) from strain 92517 *Nocardiopsis* showed the highest productivity with a peak intensity of 4.59E8, ten time more than the other three extracts.
ù

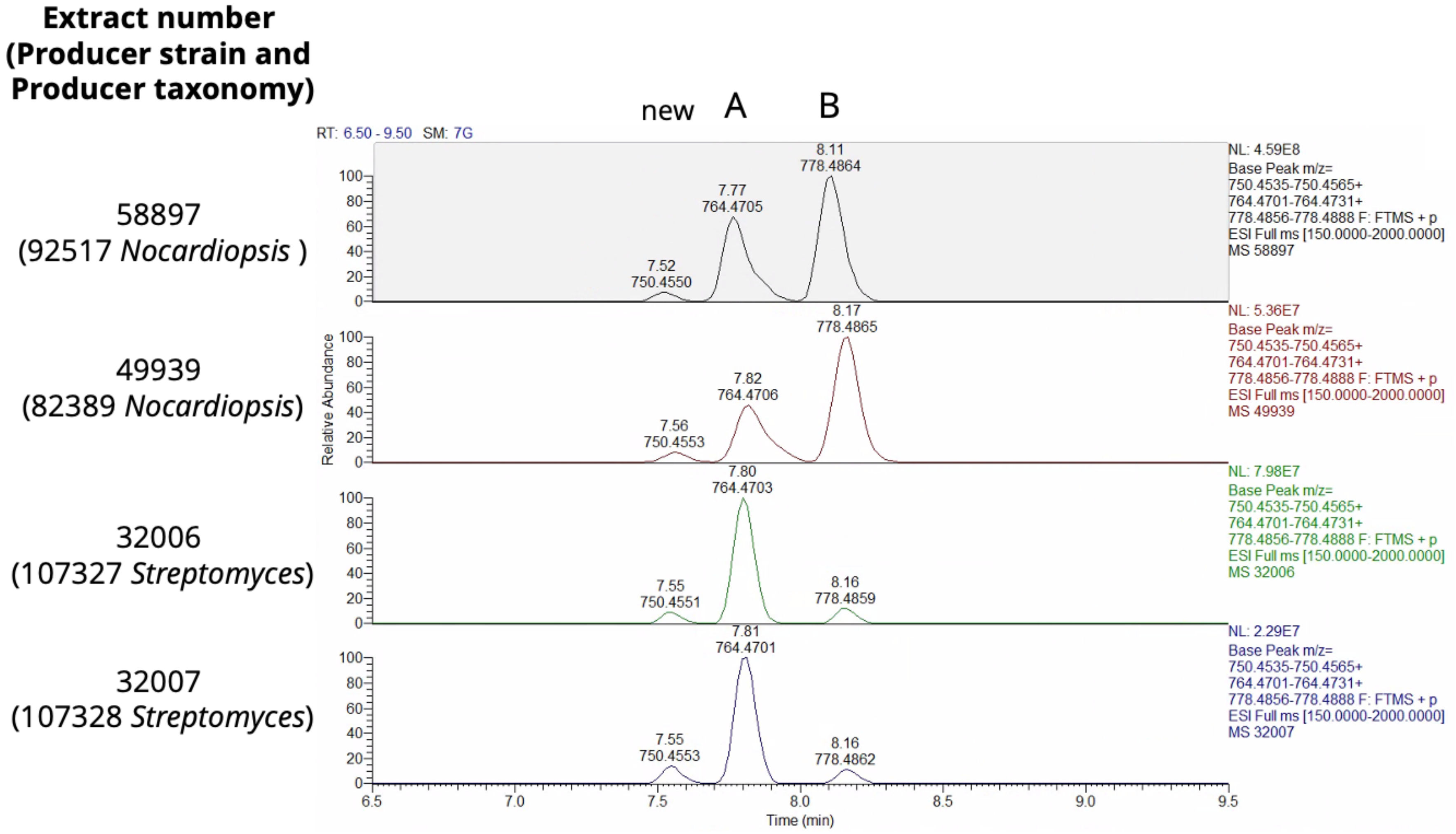

**Supplementary Figure 7:** Scatter plot correlating Retention time and exact mass for the 58,093 molecules annotated at the 1–5 confidence levels.

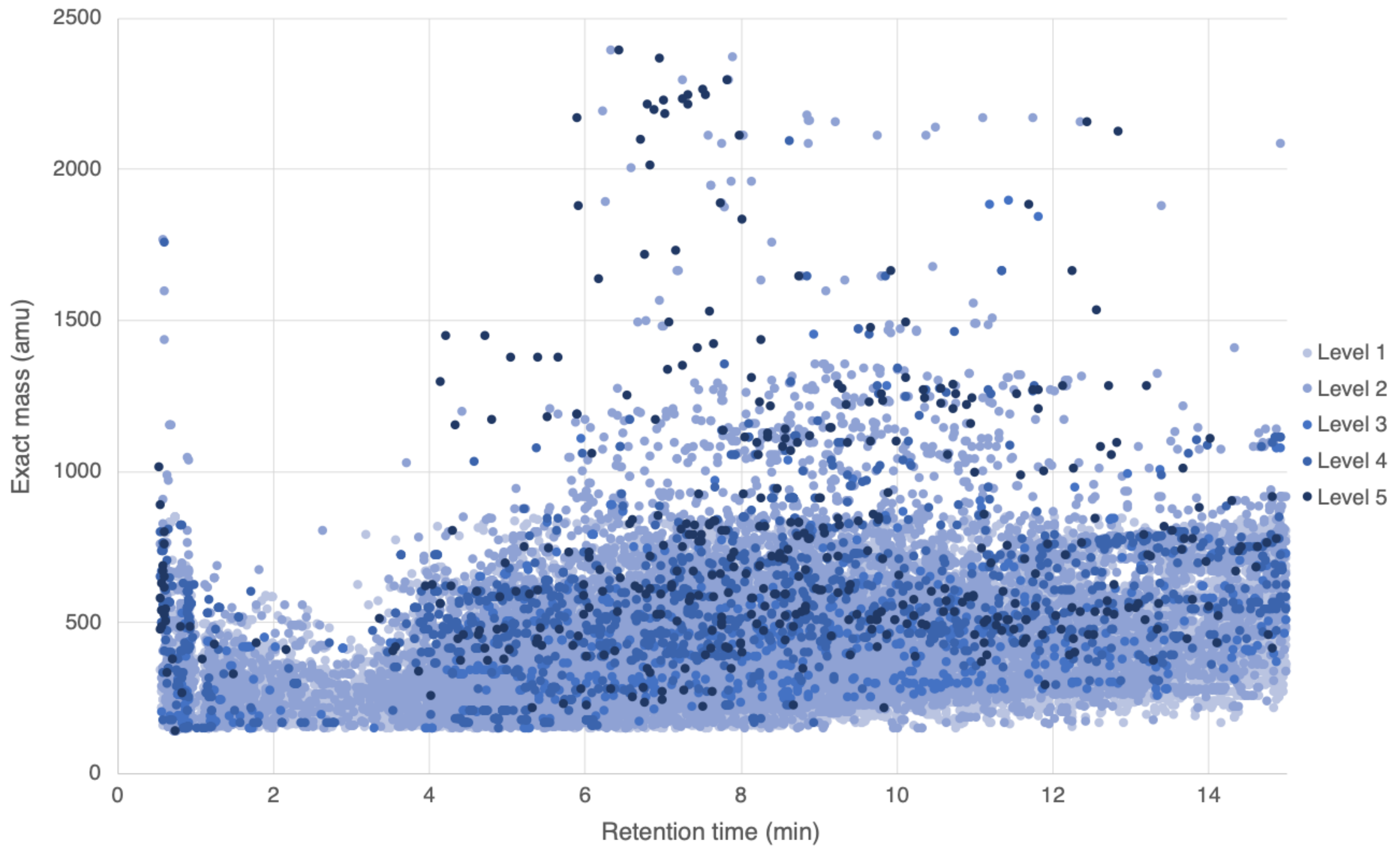
